# m^6^A-dependent microRNA binding to chromatin-associated RNA for transcriptional activation

**DOI:** 10.64898/2026.02.02.703428

**Authors:** Yuhao Zhong, Lei Zheng, Jiahao Li, Chang Liu, Jiangbo Wei, Chang Ye, Xiaoyang Dou, Bei Liu, Elias Barbosa, Fan Yang, Sean P. Pitroda, Mengjie Chen, Ralph R. Weichselbaum, Chuan He

## Abstract

For decades, microRNAs (miRNAs) have been canonically viewed as post-transcriptional repressors. We discovered extensive binding of microRNAs to chromatin-associated RNAs (caRNAs) and uncovered an *N*^6^-methyladenosine (m^6^A)–dependent transcriptional activation mechanism of microRNAs. We show that m^6^A-binding proteins FXR1/2 anchor AGO1/2 at m^6^A-marked caRNAs, where AAGUGC-seed microRNAs function as guide RNAs to direct AGO positioning. This dual anchoring stabilizes the AGO-microRNA/FXR-m^6^A complex at specific loci, which in turn recruits the ATP-dependent chromatin remodeler SMARCA4 (BRG1) to promote local chromatin opening and TET1 for DNA demethylation, respectively. Together, these coordinated activities establish a transcriptionally permissive chromatin environment, enhancing accessibility and transcription across hundreds of genes in diverse cell types. Beyond the AAGUGC-seed family, additional microRNAs and siRNAs also enhance transcription, suggesting that caRNA binding and transcriptional activation may represent a broader property of small RNAs.

## Introduction

MicroRNAs (miRNAs) are endogenous ∼22-nucleotide RNAs, initially discovered in C. elegans that associate with Argonaute (AGO) proteins to repress gene expression post-transcriptionally through mRNA degradation or translational repression (D. P. Bartel, 2004; W. Filipowicz, S. N. Bhattacharyya, N. Sonenberg, 2008; D. P. Bartel, 2009; R. C. Lee, R. L. Feinbaum, V. Ambros, 1993; B. J. Reinhart et al., 2000; J. Liu et al., 2004; F. V. Rivas et al., 2005; L. P. Lim et al., 2005; D. Baek et al., 2008; M. Selbach et al., 2008; H. Guo, N. T. Ingolia, J. S. Weissman, D. P. Bartel, 2010). This canonical, repressive framework places microRNAs almost exclusively in the cytoplasm. However, microRNAs biogenesis begins in the nucleus (Y. Lee, K. Jeon, J. T. Lee, S. Kim, V. N. Kim, 2002; G. Hutvagner et al., 2001; Y. Lee et al., 2003; R. Yi, Y. Qin, I. G. Macara, B. R. Cullen, 2003; R. I. Gregory et al., 2004; A. M. Denli, B. B. Tops, R. H. Plasterk, R. F. Ketting, G. J. Hannon, 2004; E. Lund, S. Guttinger, A. Calado, J. E. Dahlberg, U. Kutay, 2004; J. Han et al., 2004; J.-E. Park *et al*., 2011), and AGO proteins are also found in the nucleus, predominantly or even exclusively in some cell types (K. C. Johnson et al., 2024; F. M. Cernilogar et al., 2011; M. Ameyar-Zazoua et al., 2012; L. Sala et al., 2023). These observations suggest that microRNAs and AGO proteins may also possess nuclear functions.

Indeed, nuclear small RNAs, including microRNAs, are recognized to contribute to transcriptional repression (F. M. Cernilogar et al., 2011; M. Ameyar-Zazoua et al., 2012; L. Sala et al., 2023; K. V. Morris, S. W.-L. Chan, S. E. Jacobsen, D. J. Looney, 2004; B. A. Janowski et al., 2006; D. H. Kim, L. M. Villeneuve, K. V. Morris, J. J. Rossi, 2006; A. A. Aravin et al., 2007; J. Han, D. Kim, K. V. Morris, 2007; D. H. Kim, P. Sætrom, O. Snøve Jr., J. J. Rossi, 2008; S. T. Younger, D. R. Corey, 2011; K. Skourti-Stathaki, K. Kamieniarz-Gdula, N. J. Proudfoot, 2014; A. A. Sarshad et al., 2018). Yet multiple studies have described RNA-induced gene activation (RNAa), in which synthetically engineered small duplex RNAs base-pair directly with promoter DNA to stimulate transcription (L.-C. Li et al., 2006; B. A. Janowski et al., 2007; R. F. Place, L.-C. Li, D. Pookot, E. J. Noonan, R. Dahiya, 200 8; M. Alló et al., 2014; V. Portnoy et al., 2016; M. Shuaib et al., 2019; J. Cordero et al., 2024). These observations established that promoter-proximal targeting by small dsRNAs can, under certain conditions, activate transcription. Beyond RNAa, endogenous microRNA may also promote transcription through RNA-targeting mechanisms(Matsui, M. et al. 2013, Shuaib, M. et al. 2019). In one study, a microRNA was reported to activate gene expression by engaging promoter-associated RNAs and recruiting the MLL histone methyltransferase complex to target loci (Matsui, M. et al. 2013). However, the rules governing microRNA targeting on chromatin, the underlying molecular pathways, and the overall prevalence of such activating events remain poorly defined, largely because the chromatin-associated binding sites of endogenous microRNAs have not been systematically mapped. Thus, although microRNAs are predominantly characterized as repressors of gene expression, the mechanisms and conditions under which endogenous nuclear microRNAs may activate rather than repress gene expression remain comparatively underexplored.

Understanding different regulatory roles of microRNAs requires clarifying where and how they bind. It is therefore critical to map confident binding sites of individual microRNAs on chromatin. Notably, such information cannot be accurately inferred from sequence-based prediction alone, as additional regulatory layers beyond seed sequence complementarity are likely to contribute to targeting specificity. One such layer may involve protein–protein interactions, in which RNA-binding proteins (RBPs) may help recruit AGO proteins to particular sites(S. Kim et al., 2021). Similarly, RNA modifications, such as *N*^6^-methyladenosine (m^6^A), may act as a recruitment platform for regulatory machinery through m^6^A-binding proteins.

m^6^A, the most abundant internal RNA modification on messenger RNA (mRNA) in mammals and plants, is installed by the METTL3–METTL14 writer complex (J. A. Bokar et al., 1997; J. Liu et al., 2014; X.-L. Ping et al., 2014; X. Wang et al., 2016; Y. Yue et al., 2018; J. Wen et al., 2018) and can be reversed by demethylases such as FTO (G. Jia et al., 2011) and ALKBH5 (G. Zheng et al., 2013). Its regulatory functions are mediated by diverse m^6^A-binding proteins (readers) that direct RNA fate by recruiting downstream effectors (X. Wang et al., 2015; X. Wang et al., 2014; H. Shi et al., 2017). Beyond these post-transcriptional roles, RNA modifications such as m^6^A and 5-methylcytosine (m^5^C) on chromatin-associated RNAs (caRNAs) have been implicated in modulating chromatin state and transcriptional activity (J. Liu et al., 2020; J.-H. Lee et al., 2021; F. Xiong et al., 2021; J. Wei et al., 2022; W. Xu et al., 2022; S. Deng et al., 2022; R. Li et al., 2023; T. Sun et al., 2023; Z. Zou et al., 2024; X. Dou et al., 2023). In our previous work, we identified AGO1 and AGO2 as top-ranked m^6^A-associated proteins on chromatin (X. Dou et al., 2023), raising the possibility that m^6^A may recruit microRNA–AGO complexes to specific genomic loci to modulate transcription.

Here, we report extensive binding of chromatin-associated RNAs (caRNAs) by the AAGUGC-seed microRNAs on caRNAs. We show that caRNA binding by these microRNAs activates transcription through an m^6^A-dependent pathway. We found that m^6^A-binding proteins FXR1/2 recognize m^6^A-marked caRNAs and recruit AGO1/2 to these regions through direct protein–protein interactions. AAGUGC-seed microRNAs then guide AGO1/2 to specific caRNA sequences within this local environment. FXR1/2 recruit TET1 for DNA demethylation, while AGO1/2 engage SMARCA4 (BRG1) to promote chromatin opening. This coordinated epigenetic–epitranscriptomic crosstalk collectively drives chromatin opening and robust transcriptional activation across hundreds of genes in diverse cell types. Our findings, therefore, reveal an m^6^A-dependent regulatory logic that may be harnessed for future transcriptional upregulation using designed small RNAs.

## Result

### AAGUGC-seed microRNAs activate *YTHDF2* transcription

The m^6^A-binding YTHDF2 protein mediates the degradation of hundreds to thousands of methylated mRNAs in an m^6^A-dependent manner. It plays a critical role in acute myeloid leukemia (AML) by regulating the degradation of *TNFRSF2* and other transcripts involved in human leukemogenesis(J. Paris et al., 2019). To explore potential interplays between YTHDF2 and other noncoding factors in leukemia, we curated public CRISPR screening data in the AML cell line K562(W.-W. Liang et al., 2024) and examined expression correlations with *YTHDF2*. Among the top five essential noncoding elements identified, the host gene of the miR-17-92 cluster, *MIR17HG*, showed a strong positive correlation with *YTHDF2* expression (Extended Data Fig. 1a,b). This cluster includes miR-17, a representative member of the AAGUGC-seed (miR-17) family, previously implicated in global gene upregulation (K. Sun et al., 2023). Because the miR-17 family is composed of six canonical AAGUGC-seed microRNAs encoded across three paralogous clusters, miR-17/20a (*MIR17HG*), miR-93/106b (*MCM7*), and miR-106a/20b (MIR106a–363 cluster), we extended our analyses to the entire AAGUGC-seed family(V. Olive, I. Jiang, L. He, 2010). Consistently, *MCM7* also displayed significant positive correlation with *YTHDF2* across AML datasets, and expression of AAGUGC-seed microRNAs closely tracked with *YTHDF2* levels (Extended Data Fig. 1b). Further analysis of patient samples revealed that *YTHDF2*, *MIR17HG*, *MCM7*, and the corresponding AAGUGC-seed microRNAs are all expressed at higher levels in AML relapse compared to primary disease (Extended Data Fig. 1c-f). These coordinated increases underscore their potential contribution to disease recurrence and suggest a functional axis of AAGUGC-seed microRNAs and YTHDF2 in AML progression.

We were intrigued by this connection and proceeded to overexpress or knock down the individual members of these microRNAs in K562 cells. Surprisingly, when we overexpressed miR-17, miR-20a, miR20b, miR93, miR106a, or miR-106b, we observed an increase in *YTHDF2* transcript level, respectively, while knockdown of each microRNA reduced the level of *YTHDF2*, without affecting other *YTHDF* homologues (Fig. 1a and Extended Data Fig. 1g,h). RNA decay assays using actinomycin D revealed minimal changes in *YTHDF2* mRNA stability, suggesting these microRNAs do not regulate *YTHDF2* via the canonical post-transcriptional decay pathway (Fig. 1b).

**Figure. 1.**
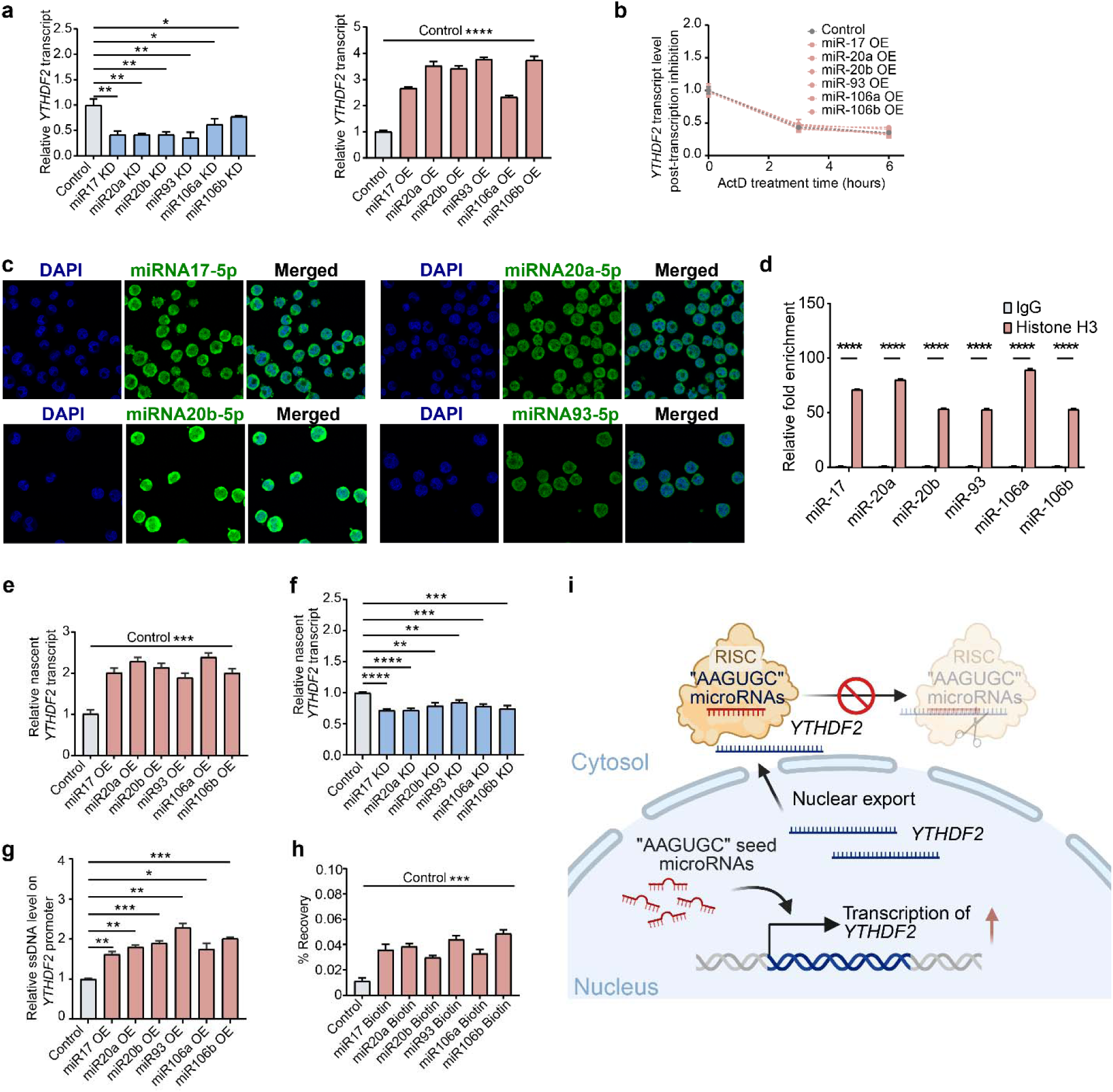
AAGUGC-seed microRNAs regulate YTHDF2 at the transcriptional level. **a**, Relative levels of *YTHDF2* transcripts quantified by qPCR in K562 cells after knockdown (left) or overexpression (right) of individual AAGUGC-seed microRNAs (miR-17, miR-20a, miR-20b, miR-93, miR-106a or miR-106b) compared with microRNA control or ASO control. **b**, *YTHDF2* transcript half-life in K562 cells overexpressing individual AAGUGC-seed microRNAs (miR-17, miR-20a, miR-20b, miR-93, miR-106a or miR-106b) compared with microRNA control. The relative levels of *YTHDF2* transcripts were normalized to the external RNA ERCC spike-in control. P-values were determined using two-way ANOVA, Dunnett’s multiple comparisons test. **c**, Representative immunofluorescence images showing nuclear localization of endogenous AAGUGC-seed microRNAs (miR-17, miR-20a, miR-20b, miR-93) in K562 cells. **d**, RNA immunoprecipitation (RIP) in K562 cells using antibodies against histone H3 or IgG, followed by qPCR quantification for mature miR-17, miR-20a, miR-20b, miR-93, miR-106a, or miR-106b. IgG is the negative control for immunoprecipitation. Data is shown as fold enrichment relative to IgG control. **e**, Relative level of nascent *YTHDF2* transcripts quantified by qPCR in K562 cells after overexpression of individual AAGUGC-seed microRNAs (miR-17, miR-20a, miR-20b, miR-93, miR-106a or miR-106b) compared with microRNA control. **f**, Relative level of nascent *YTHDF2* transcripts quantified by qPCR in K562 cells after knockdown individual AAGUGC-seed microRNAs (miR-17, miR-20a, miR-20b, miR-93, miR-106a or miR-106b) compared with ASO control. **g**, Relative ssDNA levels at the *YTHDF2* promoter in K562 cells overexpressing AAGUGC-seed microRNAs (miR-17, miR-20a, miR-20b, miR-93, miR-106a or miR-106b) compared with microRNA control. **h**, Chromatin-associated RNA immunoprecipitation (RIP) to capture transfected 3’-biotinylated microRNA mimics in K562 cells, followed by qPCR quantification of the *YTHDF2* transcript. **i**, Model of transcriptional activation of *YTHDF2* transcript by AAGUGC-seed microRNAs in the nucleus. For panels a, b, d, e, f, g, and h, p-values were determined using unpaired two-tailed Student’s t-tests (*n* = 3 biological replicates; mean ± SD shown).

We next examined the subcellular localization of these microRNAs and found them also located in the nucleus by microRNA fluorescence in situ hybridization (FISH) (Fig. 1c). Chromatin RIP-qPCR targeting histone H3 further demonstrated their association with chromatin (Fig. 1d). This nuclear and chromatin localization prompted us to investigate their potential role in transcriptional regulation. Indeed, nascent RNA synthesis assays revealed significantly elevated *YTHDF2* transcription upon miR17, miR20a, miR20b, miR93, miR106a, or miR106b overexpression (Fig. 1e), whereas microRNAs knockdown reduced *YTHDF2* transcription (Fig. 1f). A low-throughput kethoxal-assisted single-stranded DNA capture-qPCR assay further showed increased single-stranded DNA at the *YTHDF2* promoter (Fig. 1g), indicating enhanced promoter accessibility. Finally, chromatin RIP-qPCR using biotin-labeled miR17, miR20a, miR20b, miR93, miR106a, and miR106b detected direct binding to *YTHDF2* transcripts in the chromatin fraction (Fig. 1h). Together, these results revealed that these AAGUGC-seed microRNAs may directly regulate *YTHDF2* expression at the transcriptional level (Fig. 1i).

### AAGUGC-seed microRNAs promote widespread transcriptional activation

To determine whether these AAGUGC-seed microRNAs broadly affect chromatin state and transcription, we measured chromatin accessibility by DNase I–TUNEL (Fig. 2a,c and Extended Data Fig. 2a,b) and nascent transcription by quantifying 5-ethynyl uridine (5-EU) nascent RNA imaging (Fig. 2b,d and Extended Data Fig. 2c,d) and 4-thiouridine (4sU) incorporation with LC–MS/MS (Fig. 2e). We observed increased chromatin accessibility and elevated nascent RNA synthesis. We next performed the spike-in-calibrated assay for transposase-accessible chromatin sequencing (ATAC-seq) and consistently observed global increases in chromatin accessibility across K562 cells overexpressing individual AAGUGC-seed microRNAs (Fig. 2f). Such an increase in chromatin accessibility was accompanied by elevated nascent transcription of protein-coding genes (Fig. 2g). In contrast, knockdown of all AAGUGC-seed microRNAs resulted in the opposite effect, with a global decrease in nascent RNA synthesis (Fig. 2h).

**Fig. 2.**
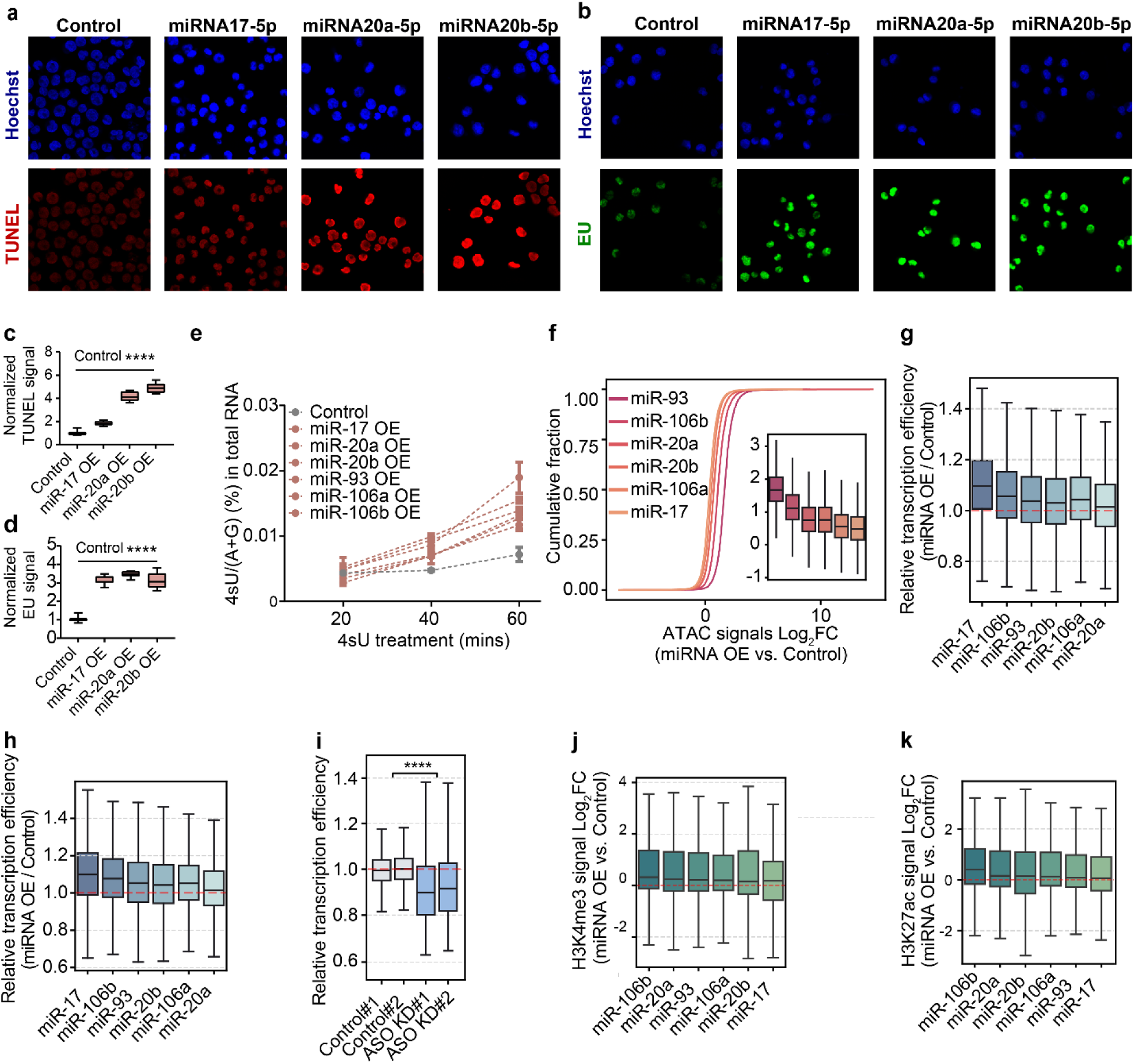
AAGUGC-seed microRNAs promote chromatin opening and overall transcription. **a**, Representative DNase I–treated TUNEL assay images (red) showing increased chromatin accessibility in K562 cells overexpressing AAGUGC-seed microRNAs (miR-17, miR-20a, or miR-20b) compared with microRNA control cells; nuclei were counterstained with Hoechst 33342 (blue). **b**, Representative EU incorporation images (green) showing increased nascent RNA synthesis in K562 cells overexpressing AAGUGC-seed microRNAs (miR-17, miR-20a, or miR-20b) compared with microRNA control cells; nuclei were counterstained with Hoechst 33342 (blue). **c**, Quantification of chromatin accessibility corresponding to the DNase I–TUNEL images in (**a**). TUNEL intensity was quantified using ImageJ from ≥5 independent biological replicates. *P*-values were determined using unpaired two-tailed t-tests. **d**, Quantification of nascent RNA synthesis corresponding to the representative EU incorporation images in (**b**). EU intensity was quantified using ImageJ from ≥5 independent biological replicates. *P*-values were determined using unpaired two-tailed t-tests. **e**, LC–MS/MS quantification of 4sU (4-thiouridine) incorporation into total RNA from K562 cells overexpressing individual AAGUGC-seed microRNAs after 20, 40 and 60□min of 4sU labeling of nascent transcripts (*n* = 3 biological replicates). **f**, Box-plots and cumulative distribution analysis of ATAC-seq log_₂_ fold-change signals (OE versus microRNA control) upon overexpression of individual AAGUGC-seed microRNAs. Each microRNA OE results in a statistically significant increase in chromatin accessibility compared with microRNA control. (\*\*\*\**p* < 0.0001, two-sided Wilcoxon’s rank-sum test). **g**, Box-plots of Relative transcription efficiency upon overexpression of individual AAGUGC-seed microRNAs. Red dashed lines indicate levels in microRNAs control. Each microRNA OE results in a statistically significant increase in nascent RNA synthesis compared with microRNA control. (\*\*\*\**p* < 0.0001, two-sided Wilcoxon’s rank-sum test). **h**, Box-plots showing log_₂_ fold changes in nascent promoter-associated RNA (paRNA) synthesis in K562 cells overexpressing individual AAGUGC-seed microRNAs (miR-17, miR-20a, miR-20b, miR-93, miR-106a or miR-106b) compared with microRNA control. Each microRNA OE results in a statistically significant increase in nascent RNA synthesis compared with microRNA control. (\*\*\*\**p* < 0.0001, two-sided Wilcoxon rank-sum test). **i**, Box-plots of relative transcription efficiency of pre-mRNA upon knockdown of all AAGUGC-seed microRNAs significantly reduces transcription efficiency compared with ASO control (\*\*\*\**p* < 0.0001, two-sided Wilcoxon’s rank-sum test). **j**, Box-plots showing log_₂_ fold changes in H3K4me3 signals in K562 cells overexpressing individual AAGUGC-seed microRNAs (miR-17, miR-20a, miR-20b, miR-93, miR-106a or miR-106b) compared with microRNA control. Each microRNA OE results in a statistically significant increase in H3K4me3 and H3K27ac signals compared with control. (\*\*\*\**p* < 0.0001, two-sided Wilcoxon rank-sum test). **k**. Box-plots showing log_₂_ fold changes in H3K27ac signals in K562 cells overexpressing individual AAGUGC-seed microRNAs (miR-17, miR-20a, miR-20b, miR-93, miR-106a or miR-106b) compared with microRNA control. Each microRNA OE results in a statistically significant increase in H3K4me3 and H3K27ac signals com pared with control. (\*\*\*\**p* < 0.0001, two-sided Wilcoxon rank-sum test).

We also observed a widespread increase in the levels of these chromatin-associated RNAs (caRNAs) in microRNA-overexpressing cells (Fig. 2i). Consistently, chromatin RIP-qPCR targeting different histone modifications revealed significantly stronger enrichment of these microRNAs by active histone modifications such as H3K4me3 but not repressive histone modifications such as H3K9me3 and H3K27me3 (Extended Data Fig. 2e). These changes were further validated by the cleavage under targets and tagmentation (CUT&Tag), which showed globally elevated H3K4me3 and H3K27ac signals in miR17, miR20a, miR20b, miR93, miR106a, or miR106b overexpressing cells, respectively (Fig. 2j,k). Together, these data indicate an microRNA-mediated gene activation process. To understand the underlying pathways and mechanisms, we, proceeded to identify RNA-binding proteins involved in this regulation.

### AGO1/2 promotes transcription and chromatin accessibility facilitated by caRNA m^6^A

The Argonaute protein family is known as the main microRNA-binding proteins that mediate transcript silencing in the cytoplasm (J. Liu et al., 2004; Rivas, F. V. et al. 2005). Because AAGUGC-seed microRNAs promote transcription and chromatin accessibility, we asked whether AGO1/2 also interacts with these microRNAs and the *YTHDF2* transcript in the nucleus. Chromatin AGO1/2 RIP-qPCR confirmed that AAGUGC-seed microRNAs and *YTHDF2* transcript are enriched in AGO1/2–bound chromatin fractions (Extended Data Fig. 3a,b). Additionally, transcript levels of *AGO1* and *AGO2* display positive correlation with that of *YTHDF2* across AML datasets (Extended Data Fig. 3c). Together, these observations indicate that AGO1/2 and AAGUGC-seed microRNAs physically associate on chromatin, suggesting a nuclear role for AGO1/2.

We further examined whether AGO1 and AGO2 depletion may affect chromatin state and transcription at their binding sites. Using low-throughput assays, we first measured nascent transcription by quantifying 4sU incorporation using LC–MS/MS. *AGO1/2* knockdown decreased 4sU incorporation into nascent RNA, indicating slower transcription (Extended Data Fig. 3d). These findings were corroborated by high-throughput sequencing, which revealed a reduction in chromatin-associated RNA expression and nascent RNA synthesis in AGO1/2-depleted cells (Fig. 3a,b).

**Fig. 3.**
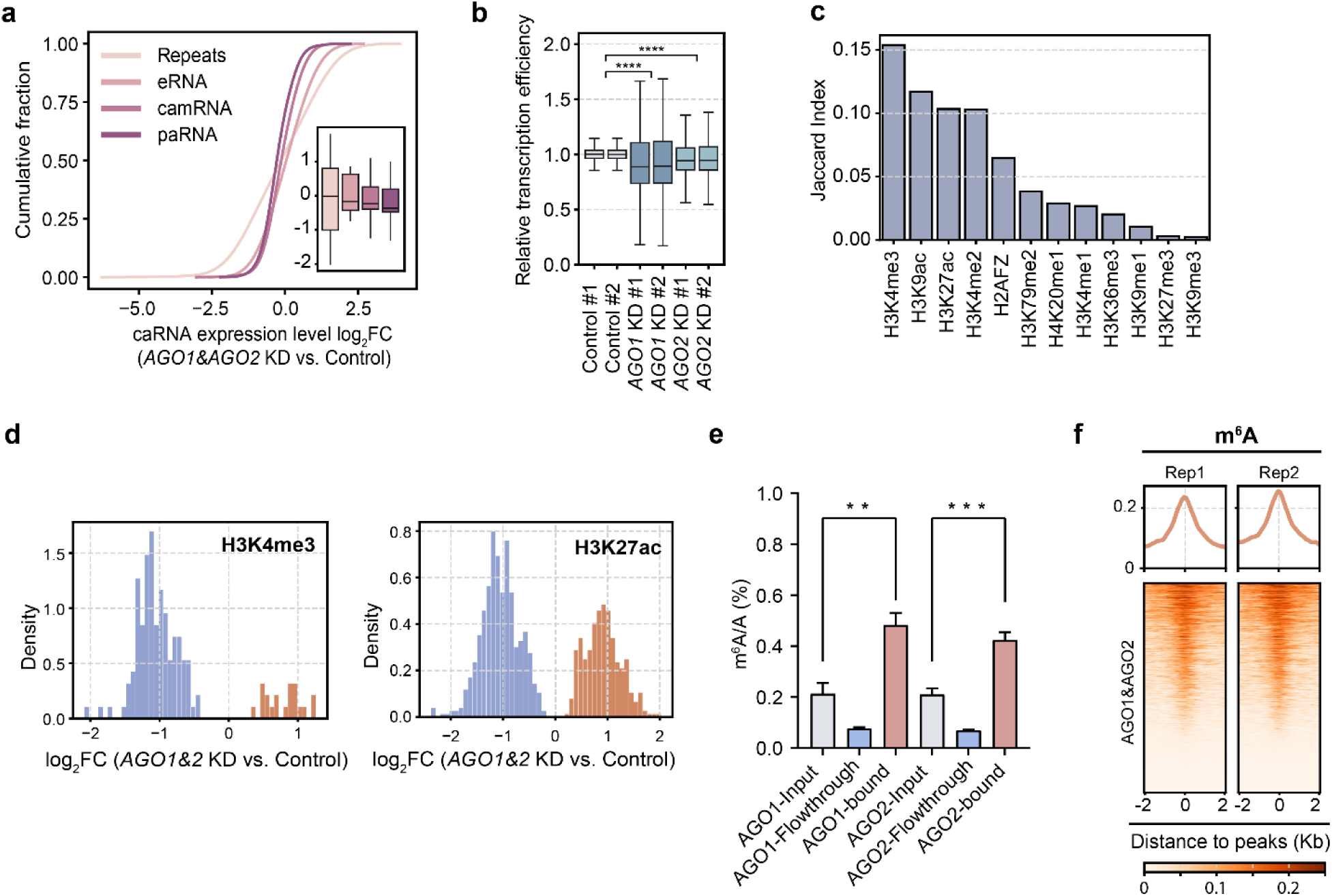
AGO1/2 promote chromatin opening and overall transcription through paRNA m^6^A. **a**, Cumulative fraction plots of log_₂_ fold change in chromatin-associated RNA (caRNA) upon *AGO1/2* KD compared with siRNA control; box-plot (right) shows distribution across categories including repeats, enhancer RNA (eRNA), chromatin-associated mRNA (camRNA), and promoter-associated RNA (paRNA). The definition and annotation of caRNA categories are provided in the Supplementary method section. *AGO1/2* KD showed a statistically significant decrease in chromatin-associated mRNA (camRNA), and promoter-associated RNA (paRNA). (\*\*\*\**p* < 0.0001, two-sided Wilcoxon’s rank-sum test). **b**, Box-plots of relative transcription efficiency of chromatin-associated pre-mRNA and paRNA upon *AGO1/2* KD. **c**, Overlap ratio of AGO1/2 peaks with histone modification ChIP–seq peaks across ENCODE datasets in K562 cells. **d**, Count of significantly different H3K4me3 (left) and H3K27ac (right) peaks in *AGO1/2* KD versus siRNA KD control K562 cells. **e**, LC–MS/MS quantification of m^6^A level in RNAs bound by AGO1, AGO2 in K562 cells. *P*-values were determined using unpaired two-tailed Student’s t-tests (*n* = 3 biological replicates; mean ± SD shown). **f**, Heatmaps and metagene profiles showing enrichment of AGO1/2 peaks around m^6^A sites in K562 cells.

Computational analysis further revealed that AGO1/2 binding sites overlap extensively with active histone modifications, including H3K4me3 and H3K27ac (Fig. 3c). Consistently, Cut&Tag profiling in AGO1/2-depleted cells further confirmed decreased H3K4me3 and H3K27ac occupancy (Fig. 3d). Together, these findings indicate that AGO1/2 regulate chromatin state and activate transcription.

RNA modifications such as m^6^A and m^5^C are known to globally influence chromatin organization and transcription (J. Liu et al., 2020; J.-H. Lee et al., 2021; F. Xiong et al., 2021; J. Wei et al., 2022; W. Xu et al., 2022; S. Deng et al., 2022; R. Li et al., 2023; T. Sun et al., 2023; Z. Zou et al., 2024; X. Dou et al., 2023). From our previous integrative analysis, AGO1 and AGO2 emerged among the top m^6^A-associated proteins on chromatin (X. Dou et al., 2023), raising the possibility that they regulate transcription in an m^6^A-dependent manner by cooperating with m^6^A and m^6^A-binding proteins. To test this, we first confirmed that m^6^A-modified RNAs were significantly enriched in AGO-bound RNA pools (Fig. 3e). Consistently, AGO-bound regions showed extensive overlap with m^6^A-modified sites (Extended Data Fig. 3e). Notably, AGO1/2 exhibited stronger binding intensity at genomic loci marked by m^6^A (Fig. 3f and Extended Data Fig. 3f).

We next analyzed AGO1/2-binding sites in microRNA-overexpressing K562 cells and found that sites proximal to m^6^A modifications exhibit increased chromatin accessibility (Extended Data Fig. 3g), enhanced nascent RNA synthesis (Extended Data Fig. 3h).These results all support caRNA binding by AGO1 and AGO2, likely guided by AAGUGC-seed microRNAs, which facilitates transcription in an m^6^A-dependent manner.

### AGO1/2-microRNA binds hundreds of caRNA sites

Although AGO1- and AGO2-binding sites exhibited increased chromatin accessibility in cells overexpressing microRNAs, it remains unclear whether microRNAs themselves are directly involved in promoting local chromatin accessibility and gene transcription. Because Argonaute proteins associate with many different microRNAs, AGO binding profiles represent the combined activity of all bound microRNAs and do not distinguish the contribution of any single microRNA. Mechanistically, how small RNAs induce gene activation has been a long-standing puzzle, in large part due to the lack of their nucleic acid binding site information. It is therefore critical to map the highly confident binding sites of these individual microRNAs on chromatin.

Conventional approaches to map microRNA binding sites like CLASH (Cross-linking, Ligation, and Sequencing of Hybrids) rely on immunoprecipitation of AGO proteins followed by proximal ligation, which can be limited by low ligation efficiency due to poor accessibility of the microRNA 3′ end (S. W. Chi, J. B. Zang, A. Mele, R. B. Darnell, 2009; M. Hafner et al., 2010; S. Grosswendt et al., 2014; M. J. Moore et al., 2015). A recently optimized ligation-based method, chimeric enhanced CrossLinking and ImmunoPrecipitation (chimeric eCLIP), combines AGO2 immunoprecipitation with simple enrichment strategies, such as PCR or biotinylated antisense oligonucleotides (ASOs) capture, that can selectively enrich for chimeric reads corresponding to specific microRNAs in the cytoplasm (S. A. Manakov et al., 2022). The low ligation and poor immunoprecipitation efficiencies, however, are still challenges when dealing with the low abundant chromatin fraction.

We comprehensively mapped the chromatin-associated RNA binding sites of AAGUGC-seed microRNAs in K562 cells using a customized method. We observed that these AAGUGC-seed microRNA-binding sites with nearby m^6^A exhibit accelerated transcription (Fig. 4a and Extended Data Fig. 4a), enhanced caRNA expression (Fig. 4b), and increased H3K4Me3 signals (Fig. 4c), upon microRNA overexpression compared with controls. These results support the direct involvement of m^6^A in this chromatin and transcription regulation pathway through these microRNAs.

**Fig. 4.**
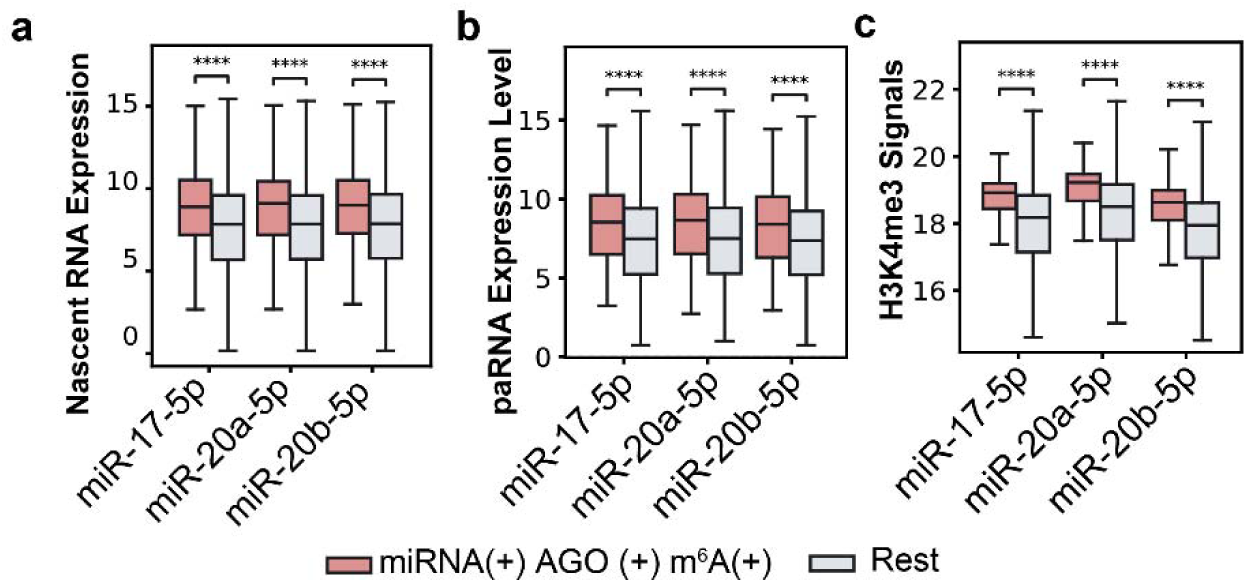
Capturing microRNAs binding sites on chromatin-associated RNAs. **a**-**c**, Box-plots comparing AGO1/2-dependent AAGUGC-seed microRNAs binding sites with nearby m^6^A versus other sites. Shown are results for miR-17, miR-20a or miR-20b overexpression, which display significantly elevated nascent RNA synthesis (**a**), paRNA expression (**b**) and H3K4me3 signals (**c**). *p*-values were determined using Wilcoxon signed-rank tests.

### AGO1/2 recruit SMARCA4 (BRG1) to promote transcription

To investigate the underlying mechanism, we studied interacting proteins of AGO1 and AGO2 and identified SMARCA4 (BRG1), the ATPase of the SWI/SNF nucleosome-remodeling complex known to promote chromatin opening and facilitate transcriptional activation. We also identified SMARCA4 (BRG1) binding as one of the top-enriched genomic features (Fig. 5a,b).

**Fig. 5.**
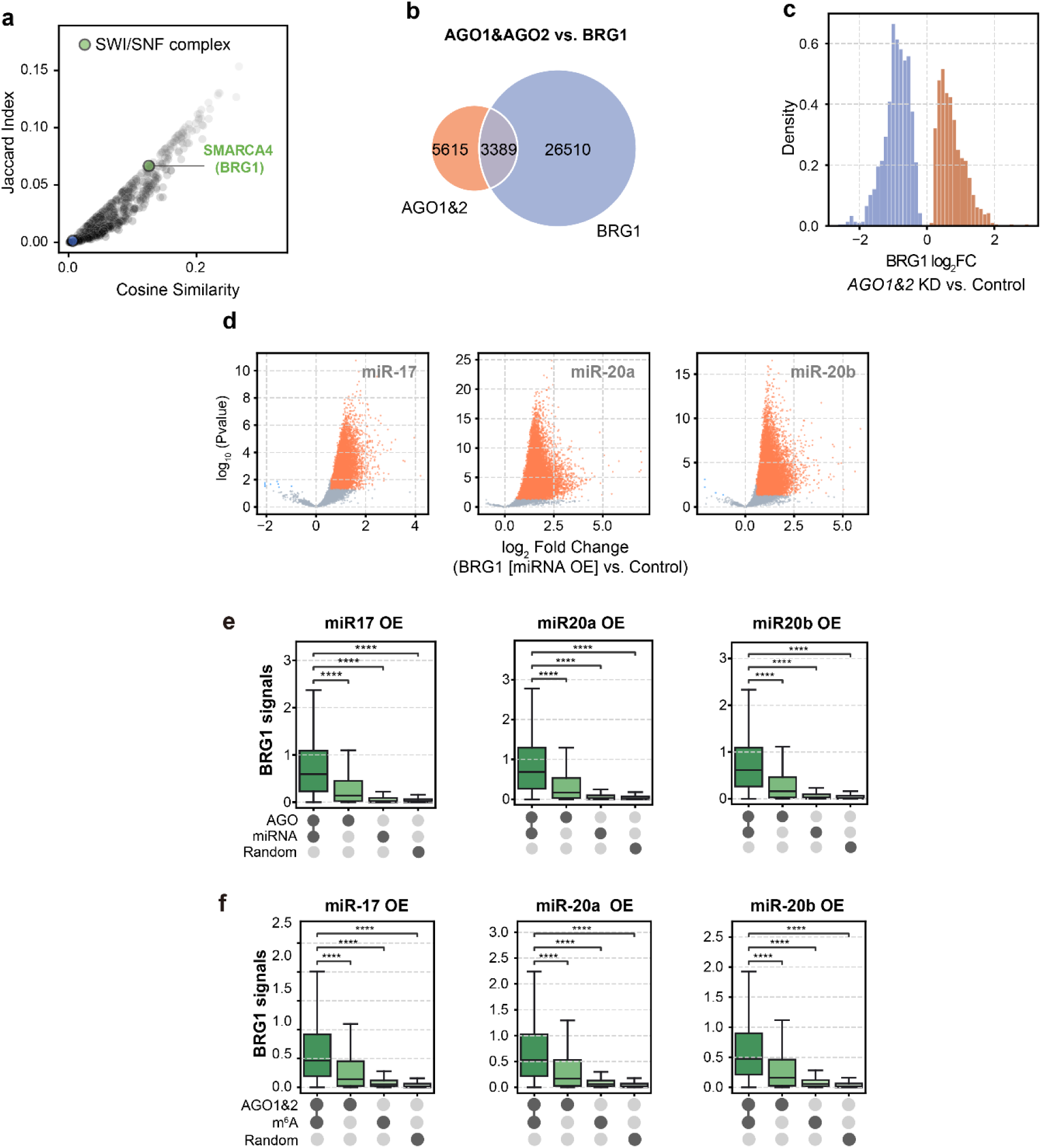
AAGUGC-seed microRNAs promote chromatin opening and activate transcription by recruiting BRG1. **a**, Jaccard index versus cosine similarity analysis of ENCODE ChIP datasets revealed strong enrichment of SWI/SNF complex binding. **b**, Overlap between AGO1/2 and SMARCA4 (BRG1) targets in K562 cells. **c**, Count of significantly different SMARCA4 (BRG1) peaks in *AGO1&2* KD versus siRNA KD control K562 cells. **d**, Volcano plots of calibrated TET1 signal changes in K562 cells overexpressing individual AAGUGC-seed microRNAs (miR-17, miR-20a, miR-20b) compared with microRNA control. **e**, Box-plots showing SMARCA4 (BRG1) signals in K562 cells overexpressing AAGUGC-seed microRNAs (miR-17, miR-20a, miR-20b). The strongest enrichment is observed at regions overlapping both AGO1&2 and AAGUGC-seed microRNAs binding sites (AGO1&2(+) miRNA(+)), whereas sites overlapping only AGO1&2 (AGO1&2(+) miRNA(−)), or only AAGUGC-seed microRNAs binding sites (AGO1&2(−) miRNA(+)) show weaker signals, and random genomic regions display minimal signals. **f**, Box-plots showing SMARCA4 (BRG1) signals in K562 cells overexpressing AAGUGC-seed microRNAs (miR-17-5p, miR-20a-5p or miR-20b-5p). The strongest enrichment is observed at regions overlapping both AGO1/2 and m^6^A peaks (+/+), whereas sites overlapping only AGO1/2 (+/−) or only m^6^A (−/+) show weaker signals, and random genomic regions (Rnd) display minimal signals.

Functionally, *SMARCA4* (*BRG1*) knockdown not only reduced nascent RNA synthesis but also reversed the transcriptional activation induced by the AAGUGC-seed microRNAs (Extended Data Fig. 5a). *AGO1&2* double knockdown diminished SMARCA4 (BRG1) recruitment, supporting that AAGUGC-seed microRNAs act via AGO proteins to engage SMARCA4 (BRG1) (Fig. 5c).

Consistent with this model, overexpression of AAGUGC-seed microRNAs enhanced SMARCA4 (BRG1) recruitment to chromatin (Fig. 5d and Extended Data Fig. 5b). Accordingly, AAGUGC-seed microRNA-binding sites and AGO1/2-binding sites proximal to m^6^A displayed stronger SMARCA4 (BRG1) occupancy in microRNA-overexpressing K562 cells (Fig. 5e,f and Extended Data Fig. 5c,d).

### The m^6^A-binding proteins FXR1/2 interact with AGO1/2 and recruit TET1

We also identified m^6^A-binding proteins such as FXR1 and FXR2. These findings support the involvement of m^6^A methylation and indicate that m^6^A-binding proteins may cooperate with the AGO–microRNA complex in this newly discovered regulation. Knockdown of either *FXR1* or *FXR2* reduced AGO1/2 RNA binding (Extended Data Fig. 6a), suggesting cooperative roles of FXR1/2 with AGO1/2. Consistently, across AML patient datasets, the expressions of *FXR1/2*, *METTL3*, *METTL14*, *DROSHA*, and *AGO1/2* positively correlated with *YTHDF2* (Extended Data Fig. 6g), supporting the functional relevance of this axis in YTHDF2 regulation.

Genome-wide mapping revealed that FXR1- and FXR2-binding sites display substantial overlaps with those of AGO1/2 (Fig. 6a). Functionally, FXR1 or FXR2 depletion impaired nascent RNA synthesis (Fig. 6b and Extended Data Fig. 6b). Additionally, FXR1 or FXR2 depletion also reduced transcriptional activation by AAGUGC-seed microRNAs (Fig. 6b and Extended Data Fig. 6b). Therefore, these results indicate that FXR1/2 can be recruited by m^6^A and AGO1/2 to these loci for transcriptional activation.

**Fig. 6.**
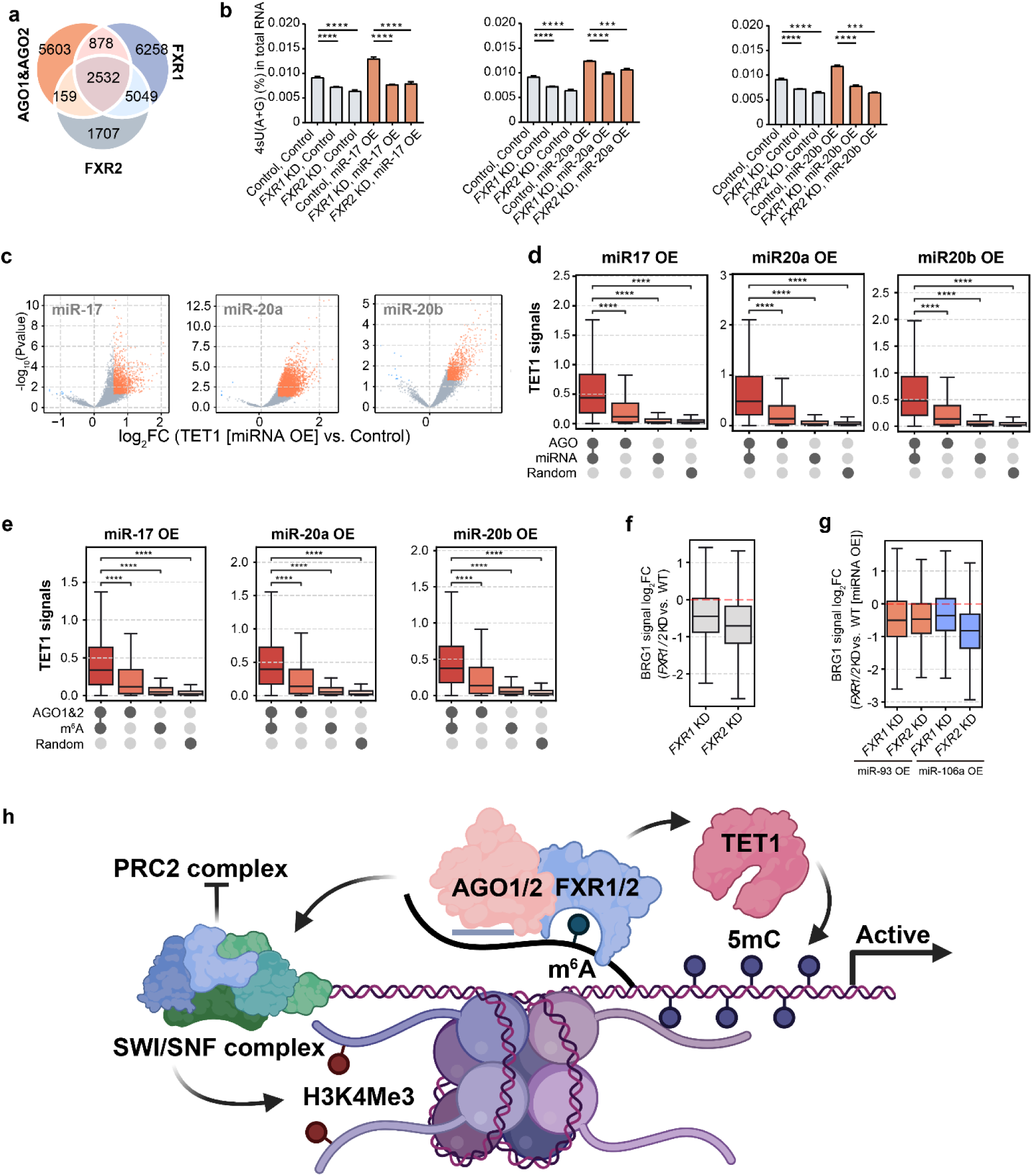
AAGUGC-seed microRNAs promote chromatin opening and activate transcription by recruiting TET1. **a**, Overlap of AGO1&2, FXR1 and FXR2 binding sites. **b**, LC–MS/MS quantification of 4sU (4-thiouridine) incorporation into total RNA after 60□min of 4sU labeling of nascent transcripts, showing that either *FXR1* or *FXR2* knockdown rescues the transcriptional activation induced by overexpression of individual AAGUGC-seed microRNAs (miR-17, miR-20a, miR-20b). *P*-values were determined using unpaired two-tailed Student’s t tests (*n* = 3 biological replicates; mean ± SD shown). **c**, Volcano plots of calibrated TET1 signal changes in K562 cells overexpressing individual AAGUGC-seed microRNAs (miR-17, miR-20a, miR-20b) compared with microRNA control. **d**, Box-plots showing TET1 signals in K562 cells overexpressing AAGUGC-seed microRNAs (miR-17, miR-20a, miR-20b). The strongest enrichment is observed at regions overlapping both AGO1&2 and AAGUGC-seed microRNAs binding sites (AGO1&2(+) miRNA(+)), whereas sites overlapping only AGO1&2 (AGO1&2(+) miRNA(−)), or only AAGUGC-seed microRNAs binding sites (AGO1&2(−) miRNA(+)) show weaker signals, and random genomic regions display minimal signals. **e**, Box-plots showing TET1 signals in K562 cells overexpressing AAGUGC-seed microRNAs (miR-17-5p, miR-20a-5p or miR-20b-5p). The strongest enrichment is observed at regions overlapping both AGO1/2 and m^6^A peaks (+/+), whereas sites overlapping only AGO1/2 (+/−) or only m^6^A (−/+) show weaker signals, and random genomic regions (Rnd) display minimal signals. (**f**–**g**) Box-plots of log_₂_ fold changes in SMARCA4 (BRG1) signal. **f**, *FXR1*/*2* knockdown versus siRNA KD control in K562 cells, showing a significant reduction in BRG1 occupancy (**** p < 0.0001, two-sided Wilcoxon’s rank-sum test). **g**, BRG1 signals in K562 cells overexpressing miR-93 or miR-106a in the *FXR1*/*2*-knockdown background, compared with their respective microRNA-overexpression condition. The microRNA-induced BRG1 increase is significantly reversed upon FXR1/2 depletion (**** p < 0.0001, two-sided Wilcoxon signed-rank test). **h**, Schematic of the proposed pathway in which AAGUGC-seed microRNAs, together with m^6^A, regulate chromatin state and activate transcription through the AGO1/2–microRNA–FXR1/2– m^6^A axis.

Mechanistically, FXR1 is known to mediate crosstalk between RNA m^6^A and DNA modification (5mC) by recruiting the DNA demethylase TET1 (S. Deng et al., 2022), thereby promoting DNA demethylation and chromatin accessibility. Consistently, we observed a high degree of overlap between FXR1/2, AGO1&2, SMARCA4 (BRG1), and TET1 binding sites, indicating frequent co-occupancy of the same genomic regions (Extended Data Fig. 6c). Moreover, overexpression of AAGUGC-seed microRNAs enhanced TET1 recruitment to chromatin (Fig. 6c and Extended Data Fig. 6d). Genome-wide distance analyses revealed that TET1 genomic binding sites displayed stronger signals at binding sites of AAGUGC-seed microRNAs and AGO1/2-binding sites proximal to m^6^A (Fig. 6d,e and Extended Data Fig. 6d,e).

FXR1/2 depletion not only reduced TET1 occupancy but also markedly decreased SMARCA4 (BRG1) recruitment to chromatin (Fig. 6f). Furthermore, the increased SMARCA4 (BRG1) occupancy induced by miR-93 or miR-106a expression was reversed upon *FXR1/2* knockdown (Fig. 6g). These findings are consistent with a model in which both FXR1/2 and AGO1/2 are required to stabilize this AGO1/2─microRNA/FXR1/2─m^6^A complex at the m^6^A-marked, microRNA-binding caRNA loci. This complex engages SMARCA4 (BRG1) through AGO1/2. SMARCA4 (BRG1), in turn, promotes deposition of active histone modifications, including H3K4me3 and H3K27ac, consistent with their globally increased signals observed in microRNA-expressing cells and decreases in AGO1/2-depleted cells. FXR1/2 recruit TET1 to promote DNA demethylation, thereby further facilitating chromatin opening and transcriptional activation. This coordinated action, anchored by two regulatory elements on caRNA, m^6^A and microRNA binding, establishes a robust transcriptionally active chromatin environment with high specificity and tenability (Fig. 6h).

### AAGUGC-seed microRNAs activate transcription in other cell types

To explore whether the transcriptional and chromatin-regulatory functions of AAGUGC-seed microRNAs extend beyond AML, we also investigated their roles in cancer and early embryonic systems. These AAGUGC-seed microRNAs tend to be upregulated the most in colon adenocarcinoma (COAD) compared to other common cancers and are known to drive cancer metastasis (Extended Data Fig. 7a) (J. Xu et al., 2019; S. P. Pitroda et al., 2018; S. J. Wang et al., 2017; G.-J. Zhang et al., 2015). Notably, miR-20a-5p, a member of this seed family, is one of only two microRNAs downregulated in patients with favorable clinical outcomes (Extended Data Fig. 7b) (A. K. Busch, T. Litman, P. S. Nielsen, 2007). Consistent with our AML findings, overexpression of AAGUGC-seed microRNAs increased *YTHDF2* expression (Extended Data Fig. 7c). Additionally, overexpression of these microRNAs enhanced transcription as indicated by increased 4sU incorporation (Extended Data Fig. 7d) and elevated chromatin-associated RNA expression transcriptome-wide in HCT116 cells (Extended Data Fig. 7e). Functionally, overexpression of these microRNAs promoted HCT116 cell proliferation and migration, effects that could be partially reversed by *YTHDF2* knockdown (Extended Data Fig. 7f,g).

AAGUGC-seed microRNAs are also essential regulators of early embryogenesis in mice. Among them, the miR-290–295 cluster is particularly dominant, with AAGUGC-seed microRNAs collectively accounting for more than 70% of the total microRNA population in mouse embryonic stem cells (mESCs) (Extended Data Fig. 7h). Overexpression of AAGUGC-seed microRNAs in mESCs induced transcriptional activation and increased chromatin accessibility patterns similar to those seen in K562 and HCT116 cells (Extended Data Fig. 7i-k). Likewise, overexpression of miR-373, another AAGUGC-seed microRNA, also increased chromatin accessibility (Extended Data Fig. 7j,k). Together, these results indicate that AAGUGC-seed microRNAs promote chromatin accessibility and transcription and act as conserved regulators of chromatin state across diverse cellular contexts.

### Transcriptional activation by non-AAGUGC microRNAs

Although our study focused on AAGUGC-seed microRNAs, we asked whether microRNA-mediated chromatin activation is restricted to this seed family. To address this question, we examined additional abundant microRNAs with distinct seed sequences. We thus generated knockout lines for multiple highly abundant microRNAs in colon cancer HCT116 cells. Loss of multiple microRNAs resulted in reduced chromatin accessibility and diminished nascent RNA synthesis, as measured by DNase I–TUNEL (Extended Data Fig. 8a,b), 4sU incorporation (Extended Data Fig. 8c), and global nascent RNA-sequencing (Extended Data Fig. 8d), suggesting that transcriptional activation may be a broader property of microRNAs.

This broader view is further supported by prior observations that transfection of diverse microRNAs and siRNAs frequently resulted in gene upregulation (Extended Data Fig. 8e) (A. Khan et al. 2029) an effect previously attributed to passive saturation of cytoplasmic AGO–RISC complexes. Consistent with this, we found that many of these previously reported microRNAs and siRNAs also increase chromatin accessibility and activate transcription in K562 cells (Extended Data Fig. 8f-h). Together, these analyses suggest an alternative interpretation of these earlier findings: that at least part of the reported global upregulation arises from transcriptional activation at chromatin and provide a potential mechanistic explanation for why microRNAs and siRNAs can induce gene upregulation across studies.

## Discussion

In this study, we uncover a distinct nuclear regulatory mechanism by which microRNAs directly regulate transcription in an m^6^A-dependent manner. We identify hundreds of endogenous microRNA-binding sites on chromatin-associated RNAs. These data reveal that a group of AAGUGC-seed microRNAs, together with Argonaute proteins, act in the nucleus to modulate chromatin state and transcription.

Our results support a model in which AAGUGC-seed microRNAs function as guide RNAs for AGO1/2 by base pairing with chromatin-associated RNAs, while m^6^A and its binding proteins FXR1/2 provide an anchoring platform that stabilizes microRNA–AGO complexes through direct protein–protein interactions. This coupling between sequence-specific RNA recognition and RNA modification–dependent anchoring enables microRNA–AGO complexes to engage chromatin-associated regulatory RNAs at defined genomic loci. Transcriptional activation occurs selectively when both a microRNA-recognition sequence and m^6^A methylation are present on chromatin-associated RNAs, and this can be modulated through perturbation of either the microRNA or the RNA m^6^A pathway components. Together, these findings expand the roles of caRNA m^6^A in chromatin and transcription regulation by establishing microRNA–AGO complexes as key co-regulators.

Although our mechanistic analyses focus on acute myeloid leukemia and AAGUGC-seed microRNAs, microRNA-mediated chromatin activation is not restricted to a single disease context or seed family. We observe similar chromatin accessibility and transcriptional activation in additional biological systems, including colorectal cancer cells and early embryonic settings, and find that other microRNAs or siRNAs with distinct seed sequences also increase chromatin accessibility and nascent transcription. These results suggest that transcriptional activation represents a more general nuclear property of microRNAs rather than a phenomenon unique to AAGUGC-seed microRNAs or to leukemia.

While microRNAs are well known to inhibit translation and promote mRNA decay, accumulating evidence indicate that small RNAs can also promote gene expression under specific contexts, including translational upregulation (Vasudevan, S., Tong, Y. & Steitz, J. A. 2007, Dai, P. et al. 2019) and transcriptional activation (L.-C. Li et al., 2006; B. A. Janowski et al., 2007; R. F. Place, L.-C. Li, D. Pookot, E. J. Noonan, R. Dahiya, 200 8; M. Alló et al., 2014; V. Portnoy et al., 2016; M. Shuaib et al., 2019; J. Cordero et al., 2024, Matsui, M. et al. 2013, Shuaib, M. et al. 2019). At the transcriptional level, small RNAs have been reported to activate gene expression through promoter-targeting mechanisms, often referred to as RNA-directed gene activation, although the underlying pathways have remained incompletely understood and, in some cases, debated. A major limitation of earlier studies has been the lack of experimentally defined binding sites for these small RNAs, particularly on chromatin-associated RNAs. By systematically mapping these interactions, our study overcomes this limitation and provides direct mechanistic insight into how endogenous microRNAs, in corporation with RNA m^6^A methylation, engage chromatin to regulate transcription.

## Acknowledgements

We are grateful to Dr. Pieter W. Faber for assistance with high-throughput sequencing and to Drs. Benjamin Stein and Samuel Weng for assistance with proteomics. We thank Drs. Alana Beadell and Chenyou Zhu for comments on the manuscript. This work was partially supported by the National Institutes of Health. We thank The Ludwig Center for Metastasis Research at the University of Chicago for partial support (S.P.P., R.R.W. and C.H.). C.H. is an Investigator of the Howard Hughes Medical Institute.

## Author contributions statement

C.H. and Y.Z. conceived the original idea and designed the initial studies. Y.Z. performed most of the experiments with assistance from J.L. and contributions from C.L., J.W., B.L., and E.B. F.Y. performed proteomic study. L.Z. performed most of the bioinformatics analyses with input from C.Y. and X.D. S.P.P. and R.R.W. contributed to colorectal cancer YTHDF2 observation. C.H. supervised the entire project. Y.Z., L.Z. and C.H. wrote the manuscript. All authors read and approved the final manuscript.

## Additional information

**Correspondence and request for materials** should be addressed to Chuan He.

## Method

### Cell culture

mES cells were maintained in DMEM supplemented with 15% heat-inactivated stem-cell-qualified FBS, 1× l-glutamine, NEAA, LIF, 1× β-mercaptoethanol, 3□µM CHIR99021, 1□µM PD0325901, and penicillin–streptomycin.

K562 cells were cultured in RPMI-1640 Medium with HEPES supplemented with 10% FBS, 2□mM l-glutamine, and penicillin–streptomycin.

HCT116 cells were cultured in McCoy’s 5A (Modified) Medium with HEPES, supplemented with 10% FBS and penicillin–streptomycin;

Drosophila S2 cells were maintained in Schneider’s Drosophila medium supplemented with 10% heat-inactivated fetal bovine serum and 1% penicillin–streptomycin at 25□°C under standard conditions

All mES, K562, and HCT116 cells were maintained at 37□°C with 5% CO_₂_.

### siRNA, microRNAs mimics, microRNAs ASO and plasmid transfection

siRNA transfections in K562 cells were performed using Lipofectamine RNAiMAX Transfection Reagent according to the manufacturer’s instructions. Transfection of microRNA mimics was carried out under the same conditions for K562, HCT116, and mES cells.

Plasmid transfections in K562 and HCT116 cells were performed using Lipofectamine 3000 Transfection Reagent according to the manufacturer’s instructions.

### microRNAs fluorescence in situ hybridization

K562 cells were seeded onto 96-well glass-bottom plates, fixed with 4% paraformaldehyde at room temperature, permeabilized with Triton X-100, and washed with DPBS. Hybridization was performed in RNA FISH Hybridization Buffer containing FAM-labeled microRNAs LNA probes, with incubation in the dark. After hybridization, cells were washed with hybridization buffer and nuclei were counterstained with Hoechst 33342. Imaging was performed using a Leica SP8 laser-scanning confocal microscope at the University of Chicago.

### Co-immunoprecipitation

K562 cells were pelleted by centrifugation at 500g for 3□min and washed twice with PBS. For co-immunoprecipitation of AGO1/2 or SMARCA4 (BRG1), The nuclear pellet was incubated in cold lysis buffer (50□mM Tris–HCl, pH 7.5, 150□mM NaCl, 1% NP40, 1:100 protease inhibitor cocktail, and 20□U□ml−1 RNase inhibitor) at 4□°C with rotation, followed by centrifugation at 15,000g for 15□min at 4□°C. Fifty microlitres of the supernatant was saved as input, and the remaining supernatant was incubated with anti-AGO1/2, anti-SMARCA4 antibodies or control IgG bound to Pierce™ Protein A/G Magnetic Beads 4□°C with rotation. Beads were washed five times with lysis buffer, and both bead-bound and input samples were mixed with 4× LDS loading buffer to a final concentration of 1× and heated before analysis by western blot.

### Nascent RNA captures and sequencing using 5EU labeling

K562 cells were seeded into 6□cm dishes at equal density in triplicate. Cells were incubated with 5-ethynyl uridine (5EU), followed by RNA extraction using TRIzol Reagent. The RNA pellet was resuspended and treated with TURBO DNase™ to remove residual DNA, then purified using the RNA Clean & Concentrator-25 kit (Zymo Research, R1017).

Drosophila 5EU-labeled spike-in RNA was added to an equal amount of total RNA from each sample. The mixture was subjected to a click reaction. Biotinylated RNA was repurified with the RNA Clean & Concentrator-25 kit (Zymo Research, R1017) and enriched using Dynabeads MyOne Streptavidin C1.

For enrichment, Dynabeads™ MyOne™ Streptavidin C1 were washed three times with 1× C1 B&W buffer (5□mM Tris, pH□7.5, 1□M NaCl, 0.5□mM EDTA, 0.2% Tween-20), then blocked in 200□µL blocking buffer (10% recombinant albumin, 1□µg/µL yeast tRNA, 1□µg/µL salmon sperm DNA in 1× C1 B&W buffer). After washing, the blocked beads were incubated with the purified RNA. The supernatant was removed, and the beads were treated with TURBO DNase™ to eliminate residual salmon sperm DNA, followed by washing in 1× C1 B&W buffer.

To elute RNA, beads were washed twice with PBS, resuspended in elution buffer (95% formamide, 10□mM EDTA pH□8.0, 1.5□mM biotin), and incubated at 70□°C for 10□min, followed by 90□°C for 5□min. The supernatant was collected and diluted into 1□ml TRIzol. RNA was purified according to the manufacturer’s instructions and resuspended in 20□µL nuclease-free water. During precipitation, 1□µL (20□µg) glycogen was added to the aqueous phase prior to isopropanol addition.

The final RNA was used for RNA-seq library preparation using the SMARTer Stranded Total RNA-Seq Kit v2 – Pico Input Mammalian. Libraries were sequenced on an Illumina NovaSeq X Plus and an Element Biosciences Aviti system (NovaX-10B-300), both generating 150□bp paired-end reads.

For low-throughput qPCR analysis, six synthetic 5EU-labeled RNA spike-in controls were used in place of the Drosophila 5EU-labeled spike-in RNA. All other steps remained unchanged.

### DNase I–TUNEL assay

K562 cells were resuspended and transferred to a 96-well glass-bottom plate prior to treatment. The DNase I–TUNEL assay was performed using the DeadEnd Fluorometric TUNEL System and Click-iT™ Plus TUNEL Assay Kits for In Situ Apoptosis Detection according to the manufacturer’s instructions with an additional DNase I treatment step. Cells were treated with DNase I before rTdT labeling. Cell nuclei were counterstained with Hoechst 33342. More than eight images were acquired per experiment using a Leica SP8 laser scanning confocal microscope at the University of Chicago. Three independent experiments were performed. The fluorescence intensity across different samples was quantified with Fiji (ImageJ) software.

### Nascent RNA imaging assay

Nascent RNA synthesis was assessed using the Click-iT RNA Alexa Fluor 488 Imaging Kit or Click-iT RNA Alexa Fluor 594 Imaging Kit according to the manufacturer’s instructions. K562 cells were incubated with 5-ethynyl uridine (5EU), then seeded onto a 96-well glass-bottom plate. Nuclei were counterstained with Hoechst 33342. Samples were imaged using a Leica SP8 laser scanning confocal microscope at the University of Chicago. Fluorescence intensity was quantified using Fiji (ImageJ). Total RNA synthesis per cell was calculated by multiplying the average fluorescence intensity by the cell area.

### CUT&Tag analysis

Cleavage under targets and tagmentation (CUT&Tag) was performed using the CUT&Tag-IT Assay Kit according to the manufacturer’s instructions. Briefly, equal amounts of Spike-In Control were added to each sample. Washed cells were bound to concanavalin A–coated beads, followed by incubation with primary antibodies against specific histone modifications. After secondary antibody binding, tagmentation was performed using a preassembled protein A–Tn5 transposase. Libraries were sequenced on an Illumina NovaSeq X Plus and an Element Biosciences Aviti system (NovaX-10B-300), both generating 150□bp paired-end reads.

### ATAC–seq analysis

ATAC–seq was performed using the TruePrep DNA Library Prep Kit V2 for Illumina according to the manufacturer’s instructions. Briefly, 50,000 cells were used per replicate, and an equal amount of Spike-In Control was added to each sample. Cells were permeabilized in buffer containing 10□mM Tris-HCl (pH 7.4), 10□mM NaCl, 3□mM MgCl_₂_, and 0.1% IGEPAL CA-630. Accessible chromatin regions were tagmented using a pre-assembled Tn5 transposome. Genomic DNA was then extracted and PCR-amplified to generate sequencing libraries. Libraries were sequenced on an Illumina NovaSeq X Plus and an Element Biosciences Aviti system (NovaX-10B-300), both generating 150□bp paired-end reads.

### Subcellular fractionation

Fractionation of K562 or HCT116 cells was performed as previously described, with an optimized NP-40 concentration. Briefly, 2.5□×□10□ to 5□×□10□ cells were collected, washed with 1□ml cold PBS, and centrifuged at 500□g for 5□min at 4□°C. The cell pellet was resuspended in 200□µl ice-cold lysis buffer (10□mM Tris-HCl, pH□7.4, 0.05% NP-40, 150□mM NaCl) by gentle flicking and incubating on ice for 5□min. Subsequently, 2.5 volumes (500□µl) of ice-cold sucrose solution (24% RNase-free sucrose in lysis buffer) were gently added to the bottom of each tube, and samples were centrifuged at 15,000□g for 10□min at 4□°C. The top 300□µl of the supernatant was collected as the cytoplasmic fraction.

The nuclear pellet was gently washed with ice-cold PBS without dislodging the pellet. Nuclei were resuspended in 100□µl pre-chilled glycerol buffer (20□mM Tris-HCl, pH□7.4, 75□mM NaCl, 0.5□mM EDTA, 0.85□mM DTT, 0.125□mM PMSF, 50% glycerol) by gentle flicking. An equal volume (100□µl) of cold nuclear lysis buffer (10□mM HEPES, pH□7.6, 1□mM DTT, 7.5□mM MgCl_₂_, 0.2□mM EDTA, 0.3□M NaCl, 1□M urea, 1% NP-40) was added, followed by two 2-second vortex pulses. Samples were immediately incubated on ice for 2□min, then centrifuged at 15,000□g for 2□min at 4□°C. The supernatant was collected as the soluble nuclear fraction (nucleoplasm).

The remaining pellet was gently rinsed with cold PBS without disturbing the pellet and collected as the chromatin-associated fraction.

To assess fractionation efficiency, 1/10 of each fraction was analyzed by Western blot using antibodies against GAPDH (cytoplasmic marker), SNRNP70 (nucleoplasmic marker), and Histone H3 (chromatin-associated marker).

### Chromatin-associated RNA-seq

The chromatin-associated fraction of K562 cells was collected as described above. The chromatin pellets were treated with TURBO DNase (Thermo Fisher Scientific, AM2238), followed by proteinase K digestion to release chromatin-associated RNA (caRNA). The purified RNA was further treated with TURBO DNase and subjected to a second round of phenol extraction to ensure complete removal of genomic DNA.

After caRNA isolation, ERCC RNA Spike-In Mix was added to each sample according to the manufacturer’s recommended ratio. RNA-seq libraries were prepared using the SMARTer Stranded Total RNA-Seq Kit v2 – Pico Input Mammalian (TaKaRa Bio, 634411) following the manufacturer’s protocol. Libraries were sequenced on an Illumina NovaSeq X Plus and an Element Biosciences Aviti system (NovaX-10B-300), both generating 150□bp paired-end reads.

### KAS-seq-qPCR

N³-kethoxal was synthesized according to a previously established protocol. An optimized KAS-seq protocol was followed to ensure efficient N³-kethoxal labeling. Cell pellets were permeabilized in ATAC Resuspension Buffer (10□mM Tris-HCl, pH□7.4, 10□mM NaCl, 3□mM MgCl_₂_) supplemented with 0.1% NP-40, 0.1% Tween-20, and 0.01% digitonin. After gentle pipetting, cells were incubated on ice, washed with 1□ml ice-cold DPBS, and centrifuged at 500□g for 5□min at 4□°C. Following an additional DPBS wash, cells were labeled with N³-kethoxal at 37□°C for 15□min with shaking at 1,300□rpm.

Genomic DNA was extracted and processed within 30□min to preserve N³-kethoxal modifications. For the click reaction, genomic DNA was suspended in 100□µL of PBS containing DBCO–PEG4–biotin in DMSO and 25□mM H_₃_BO_₃_ (pH□7.0) and incubated at 37□°C for 1.5□h with shaking at 1,300□rpm. RNase A was then added, followed by a 5□min incubation at 37□°C.

Biotinylated genomic DNA was purified using the DNA Clean & Concentrator-5 Kit (Zymo Research, D4013) and eluted in 100□µL nuclease-free water supplemented with 25□mM H_₃_BO_₃_ (pH□7.0). DNA was fragmented to 150–350□bp using a Bioruptor Pico (30□s on/30□s off, 30 cycles). One percent of the fragmented DNA was reserved as input, and the remaining 99% was incubated with pre-washed Dynabeads MyOne Streptavidin C1 in 1× C1 B&W buffer at room temperature for 30□min to enrich biotin-labeled DNA. Beads were washed five times, and bound DNA was eluted in 25□µL of nuclease-free water by heating at 95□°C for 10□min with shaking at 1,300□rpm. Eluted DNA and corresponding input controls were used for downstream qPCR analysis.

### RNA modifications level quantification by LC-MS/MS

Approximately total RNA or ribosome-depleted RNA was digested with Nucleoside Digestion Mix (NEB, M0649S) at 37□°C for 1□h. The digested sample was diluted to a final volume of 50□µL and analyzed by LC–MS/MS. Nucleosides were separated by reverse-phase ultra-performance liquid chromatography (UPLC) on a C18 column and detected using an Agilent 6460 QQQ triple-quadrupole mass spectrometer operating in positive electrospray ionization mode.

### Cell proliferation assay

HCT116 cells were seeded into 96-well plates for proliferation assays using the CellTiter 96® AQueous One Solution Cell Proliferation Assay, following the manufacturer’s instructions. MicroRNA transfection was initially performed in 6-well plates. At 8□h after transfection, cells were resuspended in complete medium, and 2,000 cells per well were seeded into 96-well plates (day 0). Cell proliferation was measured every 24□h by incubating cells with MTS reagent at 37□°C for 1□h.

### Cell migration assay

HCT116 cells were transfected as indicated and, 24 h after transfection, replated for migration assays using a 96-well, 8 µm pore size cell migration/chemotaxis kit following the manufacturer’s instructions.

### 5EU ERCC spike-in preparation

To generate DNA templates, ERCC RNA standards (Invitrogen, 4456740) were first reverse transcribed into cDNA using the PrimeScript RT Reagent Kit (Takara, RR036A). Specific ERCC fragments were then amplified by PCR with primers containing a T7 promoter sequence, yielding double-stranded DNA templates for in vitro transcription. PCR products were purified using the QIAquick PCR Purification Kit for PCR Cleanup (Qiagen, 28104) according to the manufacturer’s instructions, yielding double-stranded DNA templates for in vitro transcription. Double-stranded ERCC DNA templates containing a T7 promoter were subsequently transcribed in vitro using the HiScribe® T7 High Yield RNA Synthesis Kit (NEB, E2040S).

Transcription reactions were performed in a 20□μl volume containing UTP and 5-ethynyl UTP (5EUTP) and incubated overnight at 37□°C. Following transcription, the reaction mixture was treated with TURBO DNase™ (Thermo Fisher Scientific, AM2238) for 30□min at 37□°C to digest residual DNA templates. The RNA products were then purified using the RNA Clean & Concentrator-25 kit (Zymo Research, R1017).

### Drosophila 5EU-labeled spike-in preparation

Drosophila cells were incubated with 5-ethynyl uridine (5EU) to generate spike-in RNA with labeling levels suitable for normalization across K562 time-course samples (20, 40, and 60□min). This intermediate labeling duration was chosen to avoid excessive spike-in signal at early time points (e.g., 20□min) while ensuring sufficient signal at later time points (e.g., 60□min). Total RNA was extracted using TRIzol Reagent (Invitrogen, 15596026), treated with TURBO DNase™ (2□U/µL; Thermo Fisher Scientific, AM2238) to remove residual DNA, and purified using the RNA Clean & Concentrator-25 kit (Zymo Research, R1017).

### Biotinylation of immunoprecipitated RNAs

Biotin labeling of immunoprecipitated RNA was performed based on a previously published protocol, with a modified RNase I concentration. To account for variable recovery of AGO1/2 during immunoprecipitation, eluates were first assessed by SDS–PAGE and Coomassie Blue staining, and RNA biotinylation was performed using equalized protein inputs.

### AGO1 and AGO2 CLIP–LC–MS/MS

K562 cells were collected by centrifugation at 600□×□g for 3□min and washed once with cold Dulbecco’s PBS (DPBS). The cell pellet was resuspended in DPBS at a concentration of 10 million cells per ml. Cells were kept on ice and UV crosslinked.

The pellet was resuspended in lysis buffer (50□mM Tris-HCl pH□7.4, 100□mM NaCl, 1% NP-40 (Igepal CA-630), 0.1% SDS, 0.5% sodium deoxycholate, and 1:200 Protease Inhibitor Cocktail III). The lysate was then treated with TURBO DNase (Thermo Fisher Scientific, AM2238) at 37□°C for 1□h to digest genomic DNA and release crosslinked protein–RNA complexes. To further reduce viscosity and enhance complex release, the lysate was sequentially passed through 16G, 21G, and 25G needles. Lysates were cleared by centrifugation at maximum speed (≥16,000□×□g) for 15□min at 4□°C.

The cleared lysate was treated with RNase I at 37□°C for 5□min with gentle shaking. The RNase I concentration was carefully optimized to generate RNA fragments of approximately 1,000–2,000 nucleotides, thereby preserving long RNA fragments while reducing crosslinking of a single RNA transcript to multiple proteins. This fragmentation improved the efficiency and specificity of subsequent immunoprecipitation. An aliquot of input was saved and stored at – 80□°C.

The remaining lysate was incubated overnight at 4□°C on an end-to-end rotator with AGO antibody–conjugated protein A beads. Following incubation, the supernatant was collected, and the beads—containing immunoprecipitated AGO–RNA complexes—were washed five times with 1□ml of wash buffer (50□mM HEPES pH□7.5, 300□mM KCl, 0.05% [v/v] NP-40, 1× Halt protease and phosphatase inhibitor cocktail, 1× RNaseOUT recombinant ribonuclease inhibitor). Each wash was performed at 4□°C for 5□min with rotation.

Input, supernatant, AGO-bound fractions were treated with proteinase K at 55□°C for 2□h. RNA was then purified using the RNA Clean & Concentrator kit (Zymo Research), including a size selection step to retain RNA fragments longer than 200 nucleotides. This step was performed to exclude small RNAs—such as microRNAs, snoRNAs, and tRNAs—which are generally much shorter than 200□nt and not the focus of this analysis. The resulting purified RNA was used for downstream sample processing followed by LC-MS/MS.

### Total RNA decay assay by qPCR

K562 cells were seeded in 6-well plates and treated with actinomycin D for 6□h, 3□h, or 0□h prior to RNA harvest. At each time point, cells were directly lysed in TRIzol Reagent (Invitrogen), and the mixture was thoroughly vortexed until the pellet was fully dissolved to ensure complete lysis. Total RNA was extracted according to the manufacturer’s instructions, and mRNA was subsequently purified using the Dynabeads™ mRNA Purification Kit (Thermo Fisher Scientific, 61006). ERCC RNA spike-in controls (Invitrogen, 4456740) were added after total RNA extraction. ERCC-specific qPCR was performed in parallel and used as an external reference for normalization across samples. Three biological replicates were generated for each time point.

### RT–qPCR

To quantify transcript expression, total RNA was reverse transcribed using PrimeScript RT Master Mix (TaKaRa Bio, RR036A) with both oligo(dT) and random hexamer primers according to the manufacturer’s instructions. The resulting cDNA was analyzed by quantitative PCR (qPCR) on a LightCycler 96 system (Roche) using FastStart Essential DNA Green Master (Roche, 06402712001) and gene-specific primers. Relative expression changes were determined using the ΔΔCt method.

### Western blot

Protein samples were prepared from cells by lysis in RIPA buffer (Thermo Fisher Scientific, 89900) supplemented with 1× Halt protease and phosphatase inhibitor cocktail (Thermo Fisher Scientific, 78441) for 30 min at 4□°C with rotation. Protein concentration was determined using a NanoDrop 8000 spectrophotometer (Thermo Fisher Scientific). Equal amounts of protein were mixed with 4× loading buffer (Bio-Rad, 1610747) to a final concentration of 1×, heated at 72□°C for 15 min, and separated on 4–12% NuPAGE Bis-Tris gels (Invitrogen, NP0335BOX). Proteins were transferred to PVDF membranes (Thermo Fisher Scientific, 88585), blocked in TBST containing 3% BSA (Millipore-Sigma, A7030) for 30 min at room temperature, and incubated with primary antibodies at 4□°C overnight. Membranes were then washed three times in 1× TBST (5 min each) and incubated with HRP-conjugated secondary antibodies for 1 h at room temperature, followed by three additional washes in 1× TBST (5 min each). Protein bands were visualized using the SuperSignal West Dura Extended Duration Substrate kit (Thermo Fisher Scientific, 34075).

---

**Extended Data Fig. 1.**
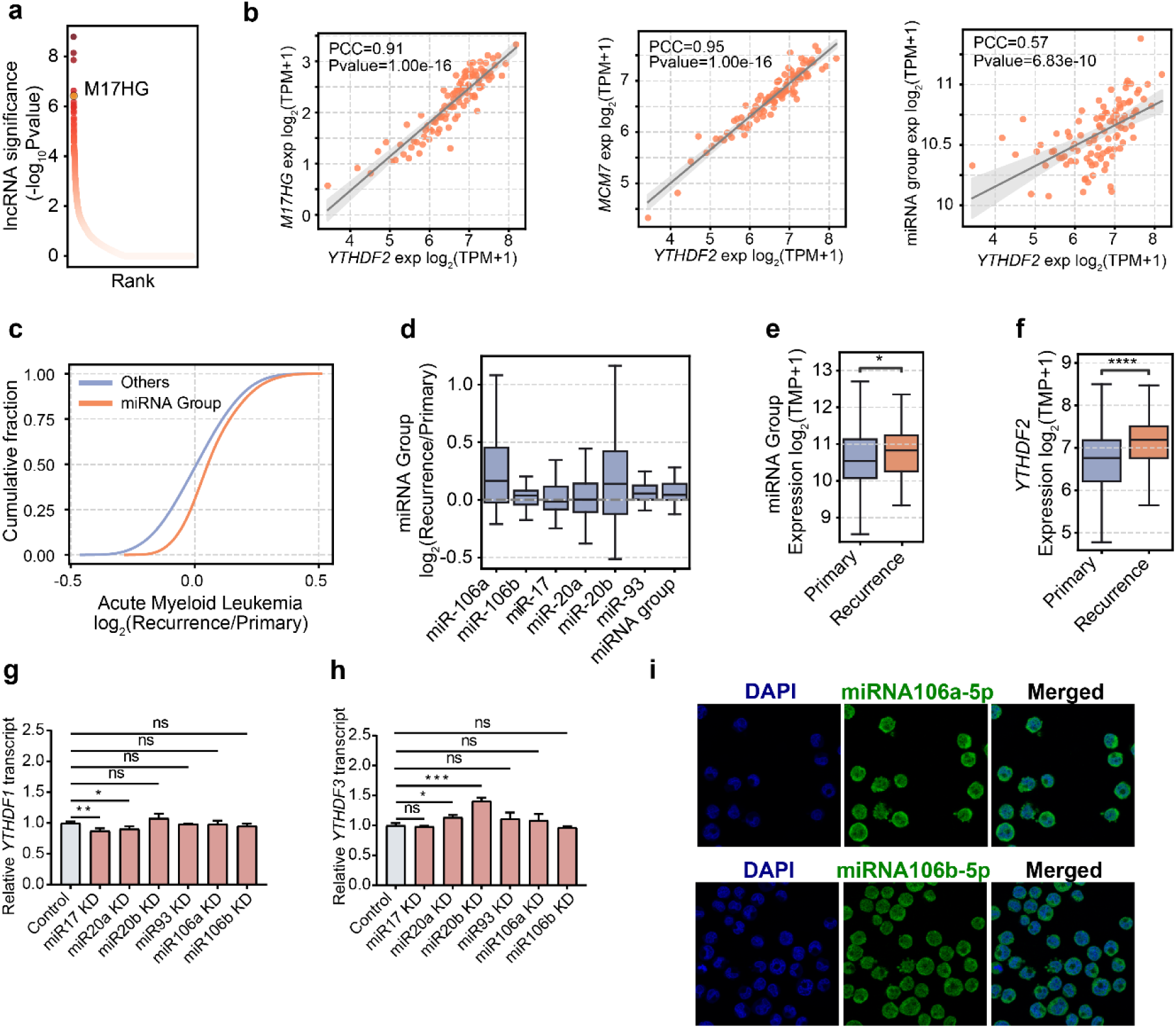
AAGUGC-seed microRNAs upregulate *YTHDF2* transcript level. **a**, Robust-rank aggregation (RRA) of lncRNAs in the K562 cells screens, based on consistent depletion of five gRNAs at 14 days after Cas13 induction. **b**, Scatter plots showing positive correlations between *YTHDF2* transcript levels and *MIR17HG*, *MCM7* and the average expression of AAGUGC-seed microRNAs (miR-17, miR-20a, miR-20b, miR-93, miR-106a and miR-106b) in the TCGA-AML cohort, with values categorized into 100 bins. PCC denotes Pearson’s correlation coefficient, and *p*-values were calculated using the t distribution. **c**, Cumulative distribution analysis of expression changes (log_₂_ recurrence/primary) in acute myeloid leukemia, comparing AAGUGC-seed microRNAs (miRNA group) with all other microRNAs (Others). **d**, Box-plots of six AAGUGC-seed microRNAs expression in primary versus recurrent acute myeloid leukemia (AML) cohorts. **e**, The average expression of AAGUGC-seed microRNAs (miR-17, miR-20a, miR-20b, miR-93, miR-106a and miR-106b) in primary versus recurrent AML samples. **f**, Expression of *YTHDF2* transcripts in primary versus recurrent AML samples. (**g**-**h**) qPCR quantification of *YTHDF1* (**g**) and *YTHDF3* (**h**) transcript levels in K562 cells after knockdown of AAGUGC-seed microRNAs (miR-17, miR-20a, miR-20b, miR-93, miR-106a or miR-106b) versus ASO control. *P*-values were determined using unpaired two-tailed Student’s t tests (*n* = 3 biological replicates; mean ± SD shown). **i**, Representative immunofluorescence images showing nuclear localization of endogenous AAGUGC-seed microRNAs (miR-17, miR-20a, miR-20b, miR-93, miR-106a or miR-106b) in K562 cells. For panels (**d**-**f**), the box plots show the median (centre line), the upper and lower quartiles (box limits) and 1.5× the interquartile range (whiskers).

**Extended Data Fig. 2.**
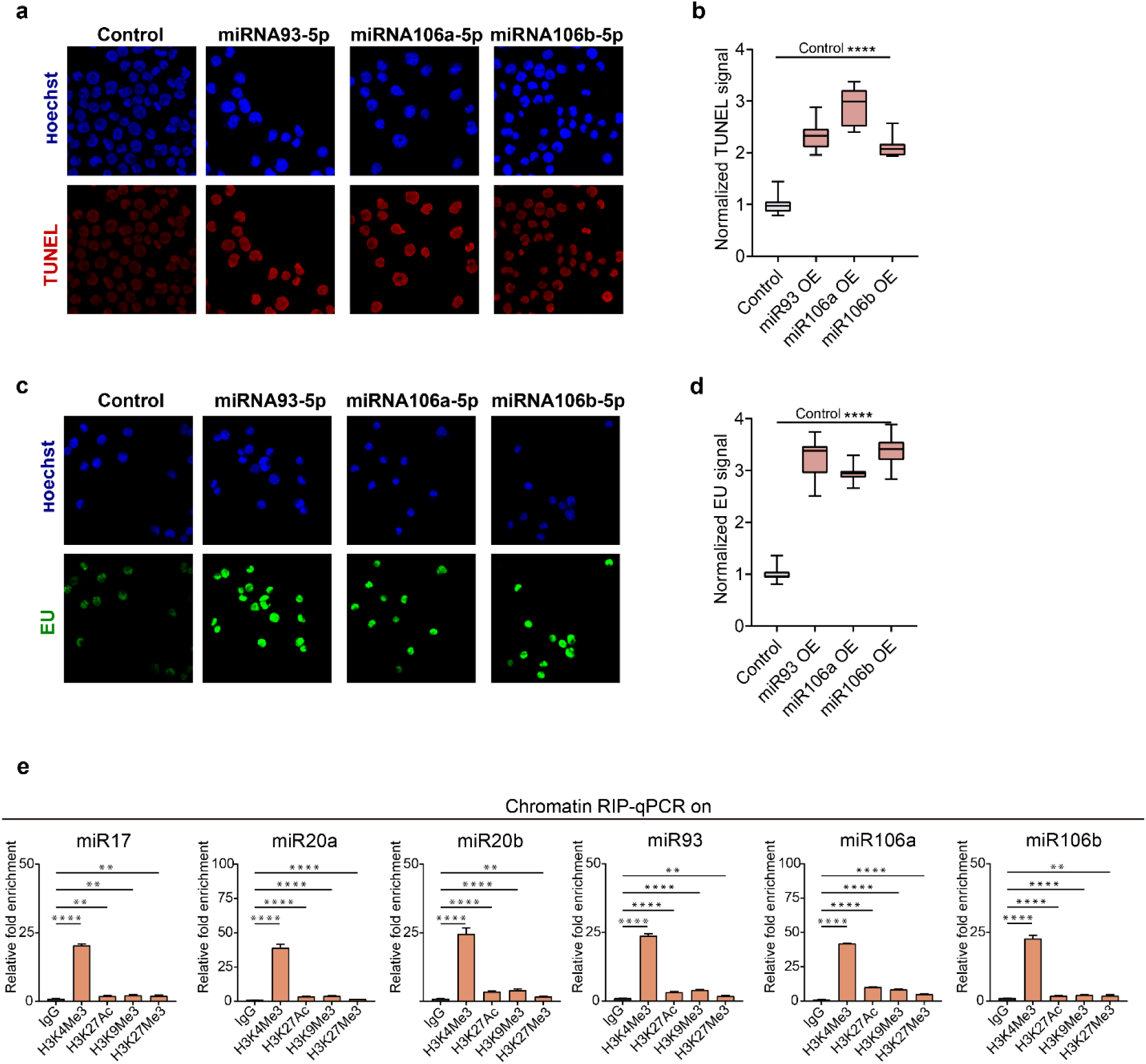
AAGUGC-seed microRNAs promote chromatin opening and overall transcription. **a**, Representative DNase I–treated TUNEL assay images (red) showing increased chromatin accessibility in K562 cells overexpressing AAGUGC-seed microRNAs (miR-93, miR-106a or miR-106b) compared with control cells; nuclei counterstained with Hoechst 33342 (blue). Quantification corresponds to data in (**b**). **b**, Box-plots showing relative DNase I-treated TUNEL assay intensity measured in K562 cells overexpressing individual AAGUGC-seed microRNAs (miR-93, miR-106a, or miR-106b); quantification corresponds to representative images shown in (**a**). TUNEL intensity was quantified using ImageJ from ≥5 independent biological replicates. *P*-values were determined using unpaired two-tailed t tests. **c**, Representative EU incorporation images (green) showing increased nascent RNA synthesis in K562 cells overexpressing AAGUGC-seed microRNAs (miR-93, miR-106a or miR-106b) compared with control cells; nuclei counterstained with Hoechst 33342 (blue). Quantification corresponds to data in (**d**). **d**, Box-plots showing relative nascent transcription intensity measured by EU incorporation in K562 cells overexpressing individual AAGUGC-seed microRNAs (miR-93, miR-106a, or miR-106b); quantification corresponds to representative images shown in (**c**). EU intensity was quantified using ImageJ from ≥5 independent biological replicates. p values were determined using unpaired two-tailed t tests. **e**, RNA immunoprecipitation (RIP) in K562 cells using antibodies against histone H3K4me3, H3K27ac, H3K9me3, and H3K27me3 or IgG control, followed by qPCR quantification for mature miR-17, miR-20a, miR-20b, miR-93, miR-106a, or miR-106b. IgG is the negative control for immunoprecipitation. Data is shown as fold enrichment relative to IgG control.

**Extended Data Fig. 3.**
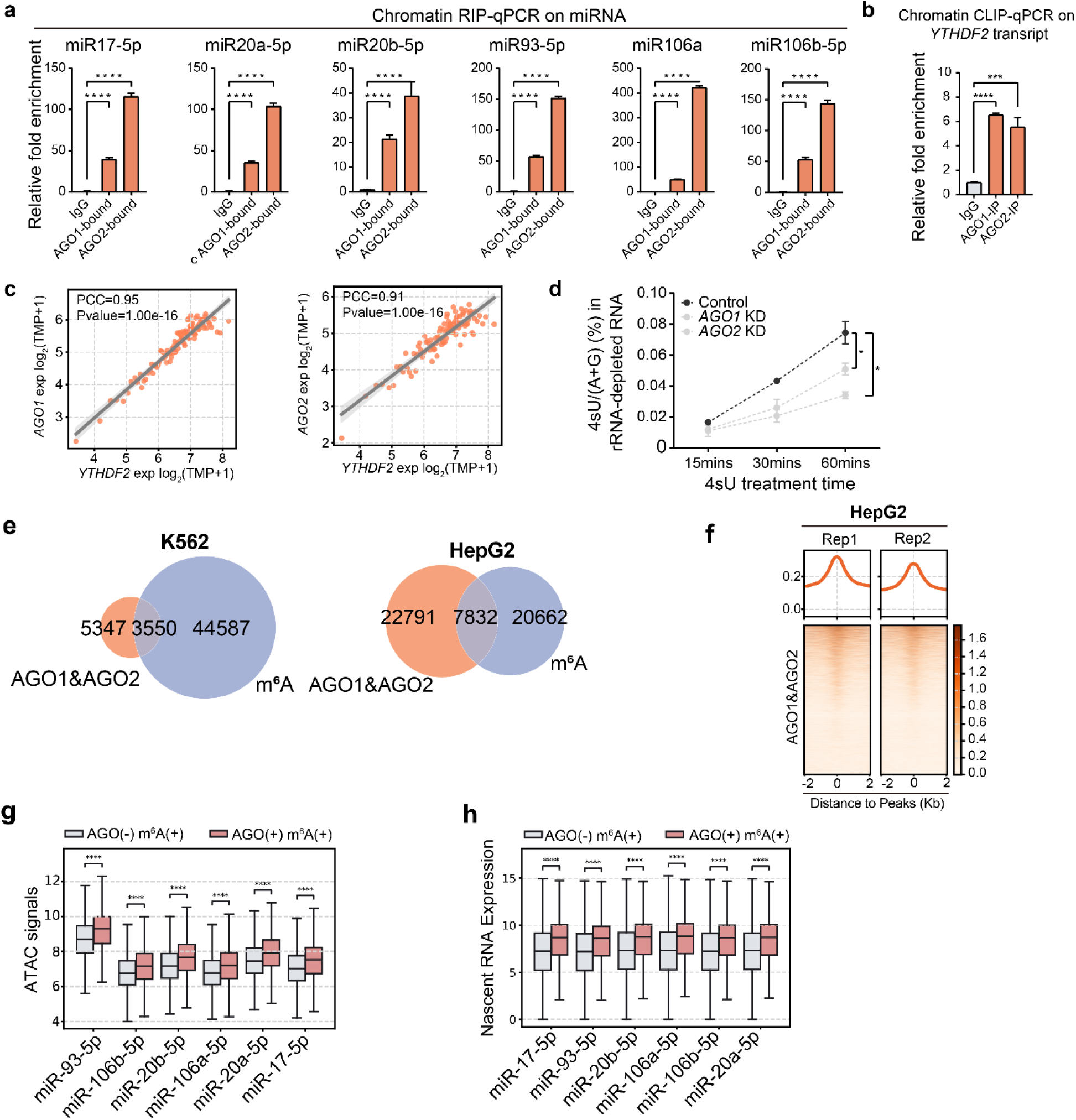
AGO1/2 may bind m^6^A-containing loci to activate transcription. **a**, Chromatin-associated RNA immunoprecipitation (RIP) in K562 cells using antibodies against AGO1, AGO2, or IgG control, followed by qPCR quantification for mature miR-17, miR-20a, miR-20b, miR-93, miR-106a, or miR-106b. Data are shown as fold enrichment relative to IgG control (*n* = 3 biological replicates). *P*-values were determined using unpaired two-tailed Student’s t tests (*n* = 3 biological replicates; mean ± SD shown). **b**, Chromatin-associated RNA immunoprecipitation (RIP) in K562 cells using antibodies against AGO1, AGO2, or IgG control, followed by qPCR quantification of *YTHDF2* transcript. Data are shown as fold enrichment relative to IgG control (*n* = 3 biological replicates). *P*-values were determined using unpaired two-tailed Student’s t tests (*n* = 3 biological replicates; mean ± SD shown). **c**, Correlation between *AGO1*, *AGO2* and *YTHDF2* expressions in the TCGA-AML cohort. Scatter plots (binned into 100 groups), showing strong positive correlations between *YTHDF2* and *AGO1* (top, Pearson correlation coefficient (PCC) = 0.95, *P* = 1.0 × 10^−16) and between *YTHDF2* and *AGO2* (bottom, PCC = 0.91, *P* = 1.0 × 10^−16). Shaded areas indicate 95% confidence intervals. **d**, LC–MS/MS quantification of 4sU (4-thiouridine) incorporation into total RNA from K562 cells with AGO1/2 depletion versus control after 15, 30 and 60□min of 4sU labeling of nascent transcripts (*n* = 3 biological replicates). *P*-values were calculated by two-way ANOVA, Dunnett’s multiple comparisons test. **e**, Overlap between AGO1&2 and m^6^A sites in K562 and HepG2 cells. **f**, Heatmaps and metagene profiles showing enrichment of AGO1&2 peaks around m^6^A sites in HepG2 cells. (**g**-**h**) Box-plots comparing m^6^A sites with nearby AGO1&2 binding sites versus m^6^A sites alone. Shown are results for overexpression of individual AAGUGC-seed microRNAs (miR-17, miR-20a, miR-20b, miR-93, miR-106a or miR-106b), which display significantly increased chromatin accessibility (**g**), elevated nascent RNA synthesis (**h**). *P*-values were determined using Wilcoxon signed-rank tests.

**Extended Data Fig. 4.**
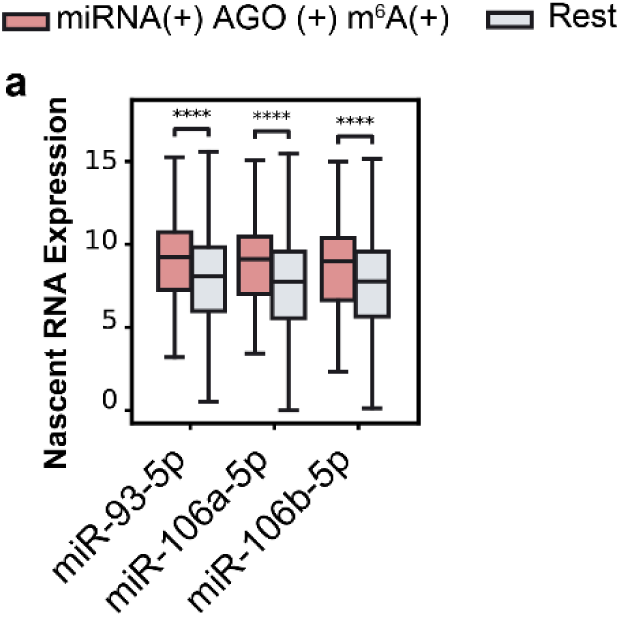
AAGUGC-seed microRNAs promote chromatin opening and activate transcription in an m^6^A-dependent manner. **a**, Box-plots comparing AGO1/2-dependent AAGUGC-seed microRNAs binding sites with nearby m^6^A versus other sites. Shown are results for overexpression of miR-93, miR-106a or miR-106b, which display significantly elevated nascent RNA synthesis. *p*-values were determined using Wilcoxon signed-rank tests.

**Extended Data Fig. 5.**
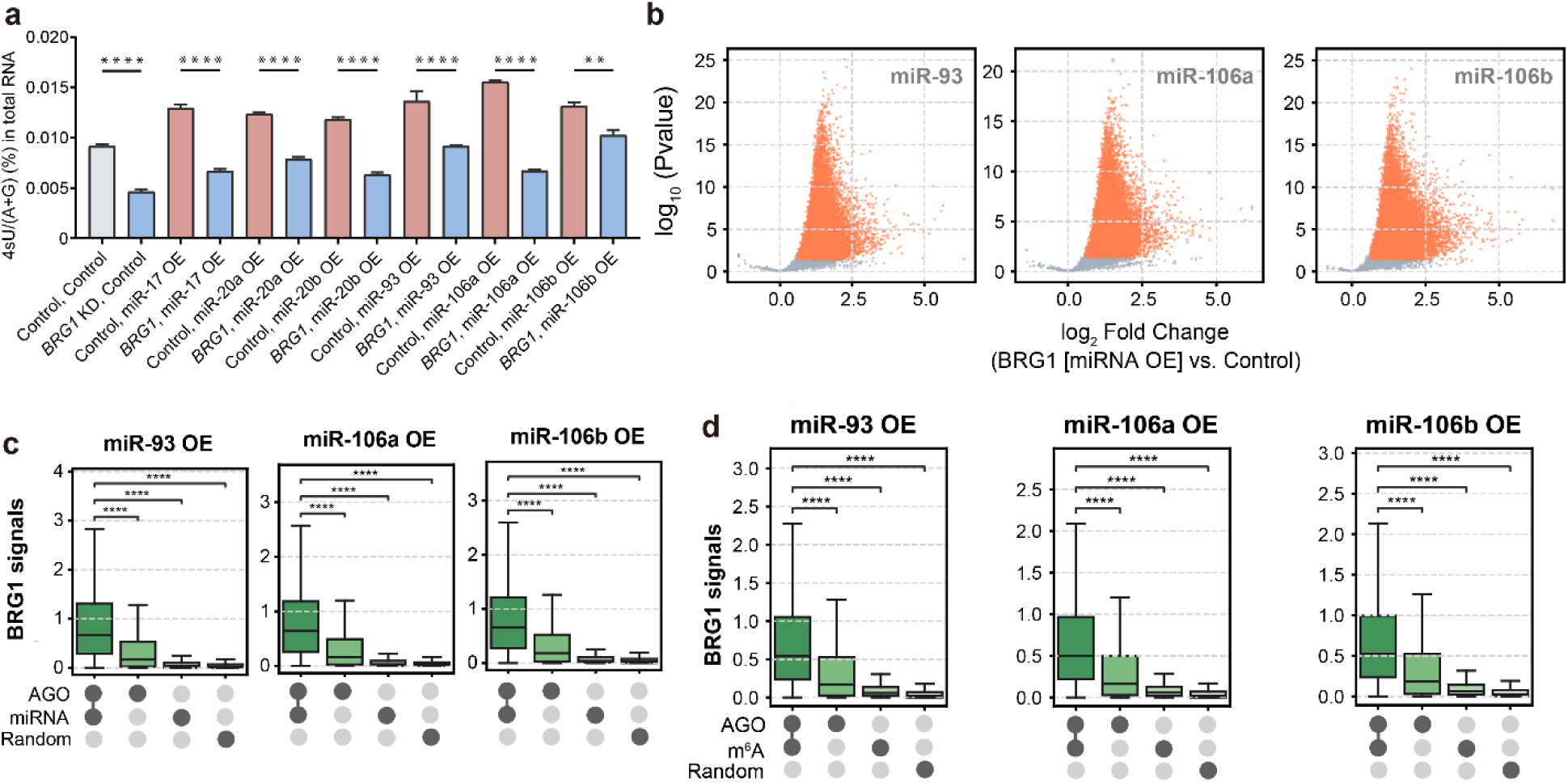
AAGUGC-seed microRNAs recruit BRG1 to nearby m^6^A loci. **a**, LC–MS/MS quantification of 4sU (4-thiouridine) incorporation into total RNA after 60□min of 4sU labeling of nascent transcripts, showing that BRG1 knockdown rescues the transcriptional activation induced by overexpression of individual AAGUGC-seed microRNAs. *P*-values were determined using unpaired two-tailed Student’s t tests (*n* = 3 biological replicates; mean ± SD shown). **b**, Volcano plots of calibrated BRG1 signal changes in K562 cells overexpressing individual AAGUGC-seed microRNAs (miR-93, miR-106a or miR-106b) compared with microRNA control. **c**, Box-plots showing SMARCA4 (BRG1) signals in K562 cells overexpressing AAGUGC-seed microRNAs (miR-93, miR-106a or miR-106b). The strongest enrichment is observed at regions overlapping both AGO1&2 and AAGUGC-seed microRNAs binding sites (AGO1&2(+) miRNA(+)), whereas sites overlapping only AGO1&2 (AGO1&2(+) miRNA(−)), or only AAGUGC-seed microRNAs binding sites (AGO1&2(−) miRNA(+)) show weaker signals, and random genomic regions display minimal signals. **d**, Box-plots showing SMARCA4 (BRG1) signals in K562 cells overexpressing AAGUGC-seed microRNAs (miR-93-5p, miR-106a-5p and miR-106b-5p). The strongest enrichment is observed at regions overlapping both AGO1&2 and m^6^A peaks (+/+), whereas sites overlapping only AGO1&2 (+/−) or only m^6^A (−/+) show weaker signals, and random genomic regions (Rnd) display minimal signals.

**Extended Data Fig. 6.**
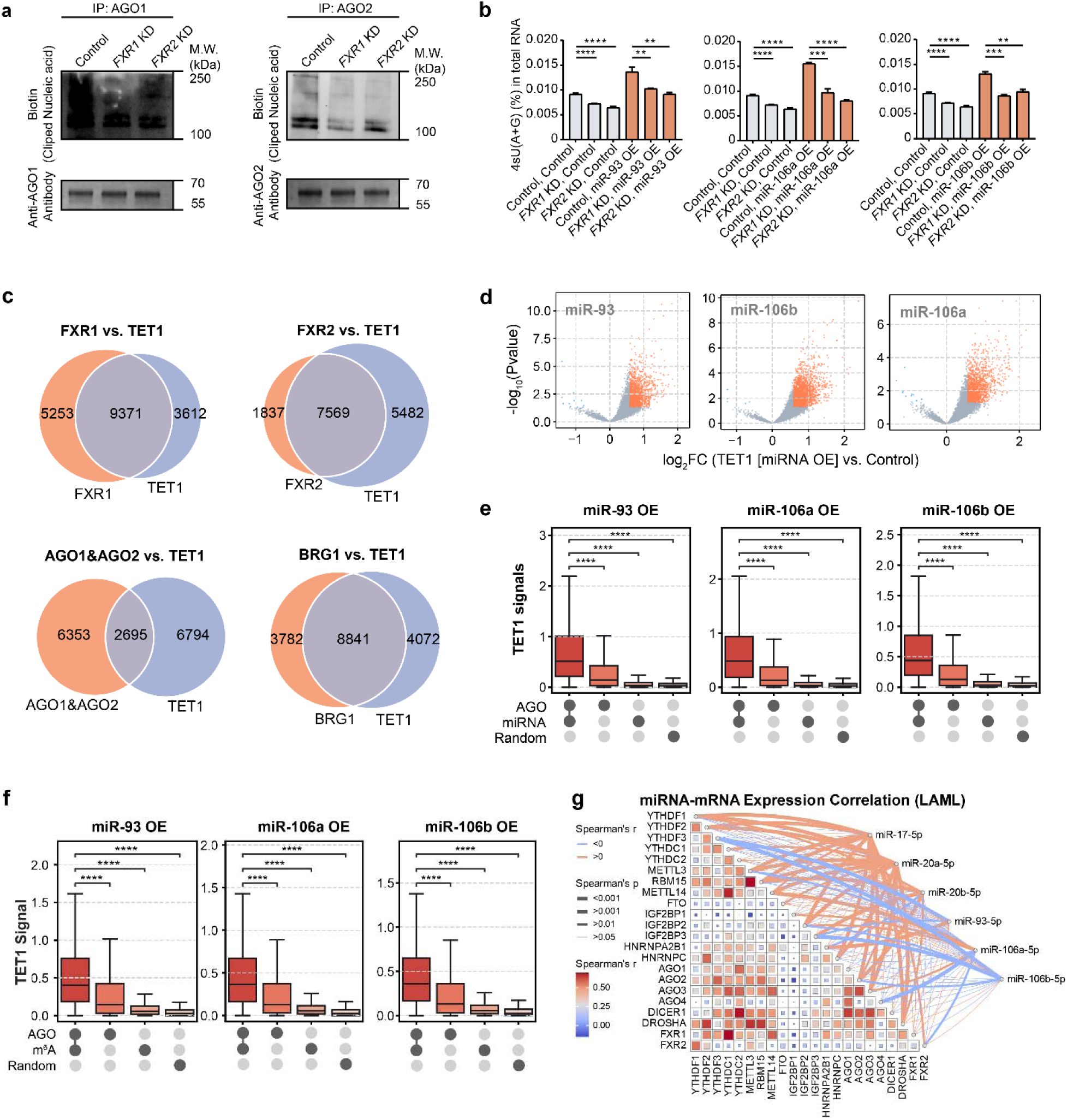
FXR1/2 mediate recruitment of AGO1/2 to nearby m^6^A loci and facilitate TET1 association. **a**, Top: western blot showing nucleic acids cross-linked to AGO1 and AGO2 in RIP samples from knockdown of either *FXR1* or *FXR2* versus control K562 cells. Bottom: Coomassie blue staining of the immunoprecipitated fractions showing IgG heavy chain, confirming equal antibody input and comparable protein loading across samples for assessment of biotin–RNA–bound proteins. **b**, LC–MS/MS quantification of 4sU (4-thiouridine) incorporation into total RNA after 60□min of 4sU labeling of nascent transcripts, showing that either *FXR1* or *FXR2* knockdown rescues the transcriptional activation induced by overexpression of individual AAGUGC-seed microRNAs. *P*-values were determined using unpaired two-tailed Student’s t tests (*n* = 3 biological replicates; mean ± SD shown). **c**, Overlap of FXR1, FXR2, AGO1&2, BRG1 and TET1 peaks in K562 cells, indicating substantial overlap with TET1-bound regions. **d**, Volcano plots of calibrated TET1 signal changes in K562 cells overexpressing individual AAGUGC-seed microRNAs (miR-93, miR-106a or miR-106b) compared with microRNA control. **e**, Box-plots showing TET1 signals in K562 cells overexpressing AAGUGC-seed microRNAs (miR-93, miR-106a or miR-106b). The strongest enrichment is observed at regions overlapping both AGO1&2 and AAGUGC-seed microRNAs binding sites (AGO1&2(+) miRNA(+)), whereas sites overlapping only AGO1&2 (AGO1&2(+) miRNA(−)), or only AAGUGC-seed microRNAs binding sites (AGO1&2(−) miRNA(+)) show weaker signals, and random genomic regions display minimal signals. **f**, Box-plots showing TET1 signals in K562 cells overexpressing AAGUGC-seed microRNAs (miR-93-5p, miR-106a-5p and miR-106b-5p). The strongest enrichment is observed at regions overlapping both AGO1&2 and m^6^A peaks (+/+), whereas sites overlapping only AGO1&2 (+/−) or only m^6^A (−/+) show weaker signals, and random genomic regions (Rnd) display minimal signals. **g**, miRNA–mRNA expression correlations in the TCGA-LAML cohort. Heatmap showing pairwise Spearman’s correlation coefficients between AAGUGC-seed microRNAs and genes in the proposed pathway, including m^6^A-binding proteins (FXR1 and FXR2), the m^6^A writer METTL3, AGO proteins and chromatin factors. Red indicates positive correlation and blue indicates negative correlation. The significance of correlations is represented by square size (larger squares denote higher *p* values). Network edges highlight significant correlations (*p* < 0.001), with red edges indicating positive and blue edges indicating negative associations. For panels (**e**) and (**f**), the box plots show the median (centre line), the upper and lower quartiles (box limits) and 1.5× the interquartile range (whiskers).

**Extended Data Fig. 7.**
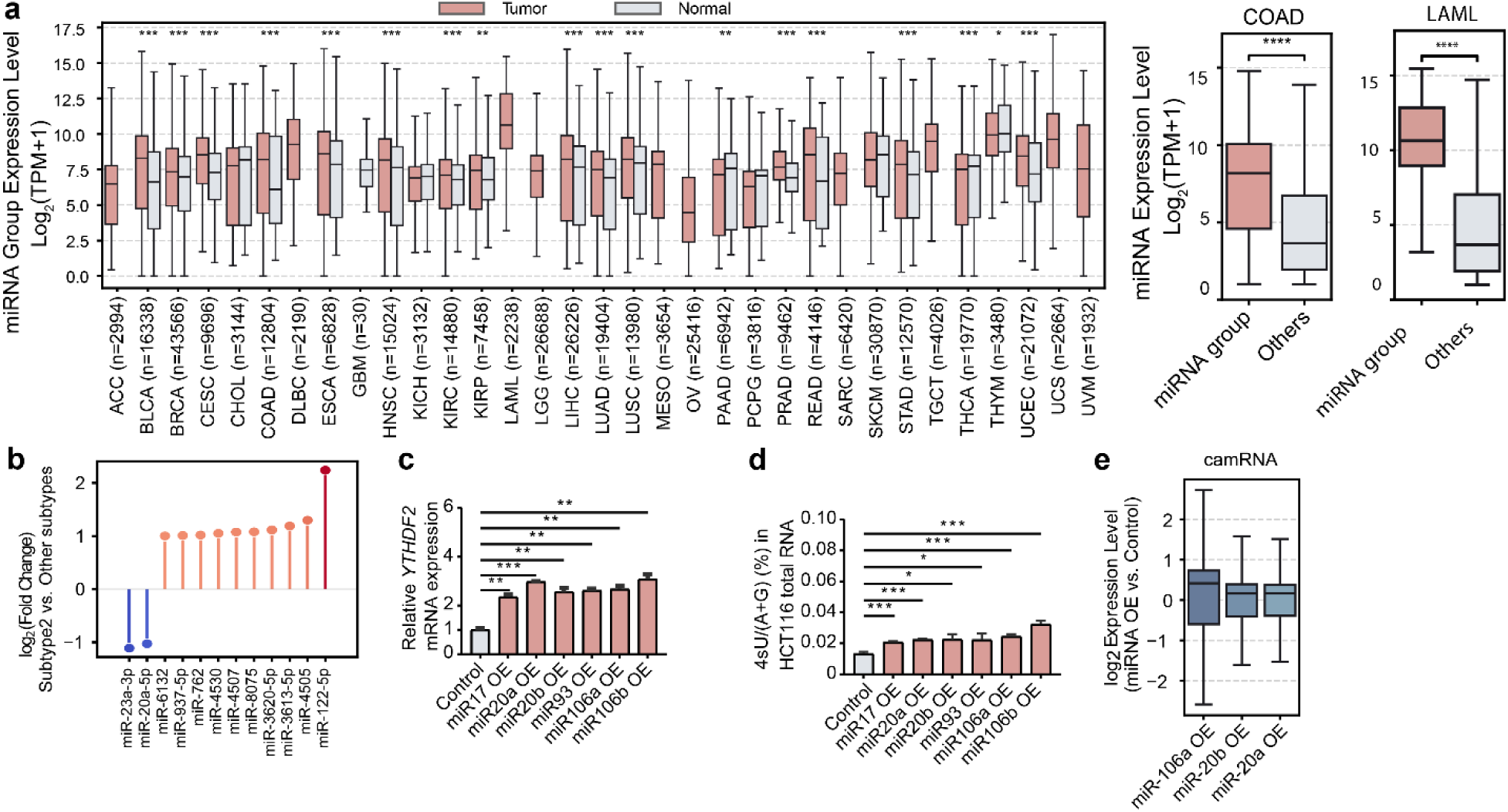
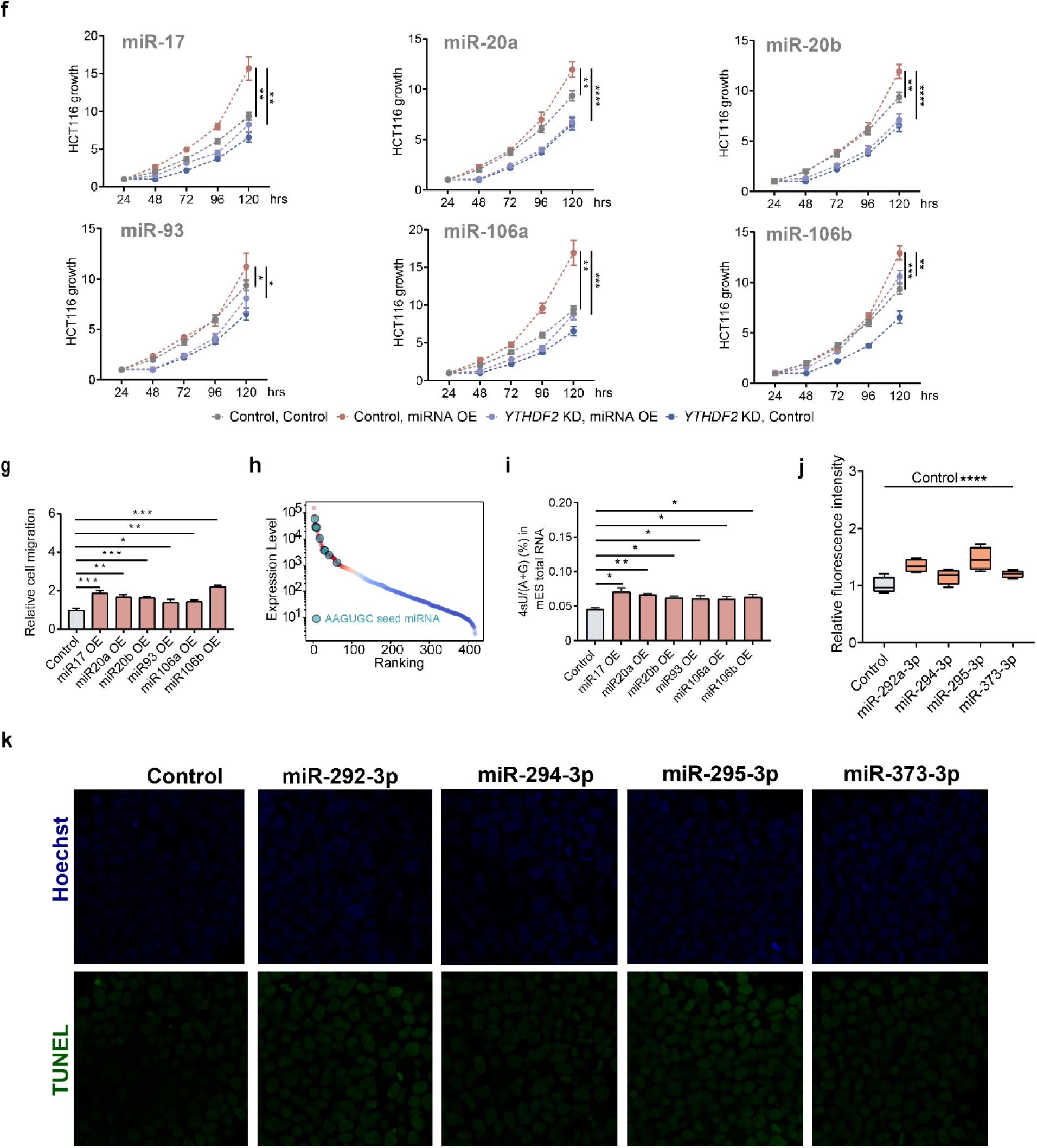
AAGUGC-seed microRNAs promote chromatin opening and overall transcription in colon cancer. **a**, Box-plots of AAGUGC-seed microRNAs expression across TCGA pan-cancer cohorts, showing significantly higher levels in tumors compared with matched normal tissues. Right, specific enrichment of the average expression of six AAGUGC-seed microRNAs (miR-17, miR-20a, miR-20b, miR-93, miR-106a and miR-106b) in colorectal adenocarcinoma (COAD) and acute myeloid leukemia (LAML). The box plots show the median (centre line), the upper and lower quartiles (box limits) and 1.5× the interquartile range (whiskers). **b**, Comparison of miRNAs expression among COAD tumor subtypes, showing reduced expression of miR-20a-5p in Subtype 2 relative to other subtypes. miR-20a-5p is one of only two miRNAs downregulated in Subtype 2, which is associated with improved clinical outcomes. **c**, Relative levels of *YTHDF2* transcripts quantified by qPCR in HCT116 cells overexpressing individual AAGUGC-seed microRNAs (miR-17, miR-20a, miR-20b, miR-93, miR-106a or miR-106b) compared with microRNA control. *P*-values were determined using unpaired two-tailed Student’s t tests. (*n* = 3 biological replicates; mean ± SD shown). *P*-values were determined using unpaired two-tailed Student’s t tests. **d**, LC–MS/MS quantification of 4sU (4-thiouridine) incorporation into total RNA from HCT116 cells overexpressing individual AAGUGC-seed microRNAs after 60□min of 4sU labeling of nascent transcripts (*n* = 3 biological replicates). **e**, Box-plots showing log_₂_ fold changes in chromatin-associated mRNA expression in HCT116 cells overexpressing individual AAGUGC-seed microRNAs (miR-20a, miR-20b or miR-106a) compared with microRNA control. Each microRNA OE results in a statistically significant increase in cam-RNA compared with microRNA control. (\*\*\*\**p* < 0.0001, two-sided Wilcoxon rank-sum test). The box plots show the median (centre line), the upper and lower quartiles (box limits) and 1.5× the interquartile range (whiskers). **f**, Proliferation curves showing that *YTHDF2* knockdown rescues the increased cell growth induced by overexpression of individual AAGUGC-seed microRNAs (miR-17, miR-20a, miR-20b, miR-93, miR-106a or miR-106b). Statistical comparisons were shown relative to the “control, miRNA OE” group at 120□h. *P*-values were calculated by two-way ANOVA, Dunnett’s multiple comparisons test. **g**, Relative migration capacity of HCT116 cells expressing individual AAGUGC-seed microRNAs (miR-17, miR-20a, miR-20b, miR-93, miR-106a or miR-106b) (*n* = 5 biological replicates). **h**, microRNA expression ranking in mouse embryonic stem cells (mES). Representative AAGUGC-seed microRNAs are highlighted. **i**, LC–MS/MS quantification of 4sU (4-thiouridine) incorporation into total RNA from mES cells overexpressing individual AAGUGC-seed microRNAs after 60□min of 4sU labeling of nascent transcripts (*n* = 3 biological replicates). **j**, Box-plots of relative DNase I-treated TUNEL assay intensity measured in (**k**). TUNEL intensity was quantified using ImageJ from ≥5 independent biological replicates. *P*-values were determined using unpaired two-tailed t tests. **k**, Representative DNase I–treated TUNEL assay images (green) showing increased chromatin accessibility in mES cells overexpressing AAGUGC-seed microRNAs (miR-292-3p, miR-294-3p, miR-295-3p, or miR-373-3p) compared with control cells; nuclei counterstained with Hoechst 33342 (blue). Quantification corresponds to data in (**j**).

**Extended Data Fig. 8.**
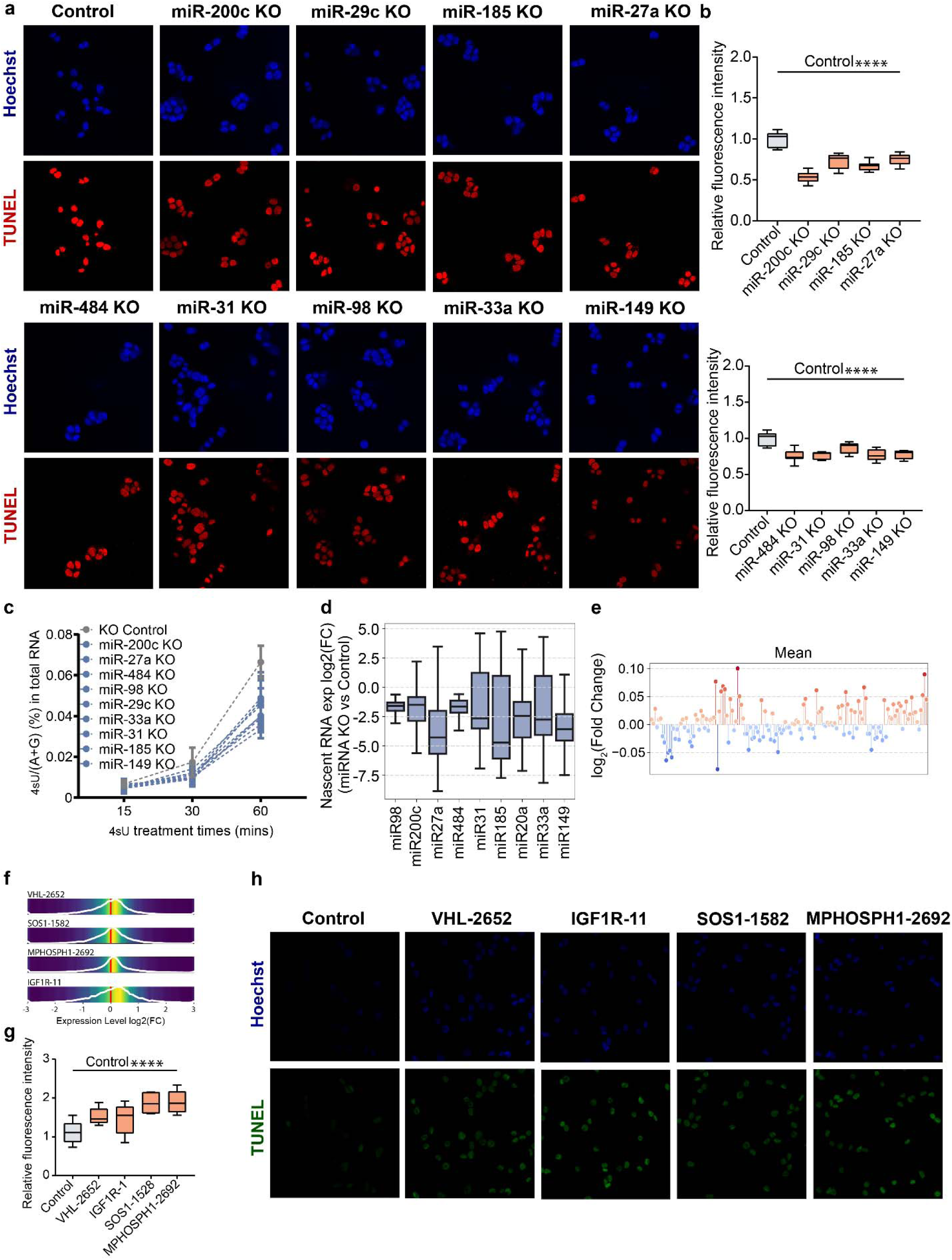
Transcriptional activation by additional microRNAs and siRNAs with distinct seed sequences. **a**, Representative DNase I–treated TUNEL assay images (red) showing increased chromatin accessibility in HCT116 cells with microRNAs knockout (KO) compared with control cells; nuclei counterstained with Hoechst 33342 (blue). Quantification corresponds to data in (**b**). **b**, Box-plots of relative DNase I-treated TUNEL assay intensity measured in (**a**). TUNEL intensity was quantified using ImageJ from ≥5 independent biological replicates. *P*-values were determined using unpaired two-tailed t tests. **c**, LC–MS/MS quantification of 4sU (4-thiouridine) levels in total RNA from HCT116 cells with miRNAs knockout after 60-minute 4sU labeling of nascent transcripts. p values were determined using unpaired two-tailed Student’s t tests (*n* = 3 biological replicates; mean ± SD shown). **d**, Box-plots showing log_₂_ fold changes in nascent RNA expression in HCT116 with microRNAs knockout (KO) compared with KO control. The box plots show the median (centre line), the upper and lower quartiles (box limits) and 1.5× the interquartile range (whiskers). All experiments and analysis were performed using two biological replicates per condition. **e**, Effect of siRNAs and microRNAs transfection on gene expression. Scatter plots showing log_₂_ fold changes across 150 RNA-seq perturbation datasets previously analyzed for individual siRNAs or microRNAs. The average gene expression of all transcripts across conditions. **f**, Distribution plots of expression changes (log_₂_ fold change) following siRNA targeting of VHL, SOS1, MPHOSPH1 and IGF1R compared with control, showing that these siRNAs induce global upregulation of gene expression. **g**, Box-plots of relative DNase I-treated TUNEL assay intensity measured in K562 cells with siRNA transfection (**h**). TUNEL intensity was quantified using ImageJ from ≥5 independent biological replicates. p values were determined using unpaired two-tailed t tests. **h**, Representative DNase I–treated TUNEL assay images (red) showing increased chromatin accessibility in K562 cells with siRNA transfection (*VHL*, *SOS1*, *MPHOSPH1* and *IGF1R*) compared with siRNA KD control cells; nuclei counterstained with Hoechst 33342 (blue). Quantification corresponds to data in (**g**).

